# Alignment-free discovery of conserved RNA structures in the human genome

**DOI:** 10.64898/2026.10.24.761493

**Authors:** Shunsuke Sumi, Koki Makino, Masahiro Onoguchi, Shungo Adachi, Hirohide Saito, Michiaki Hamada

## Abstract

Many functional RNAs were first found as conserved RNA structures, and their roles in gene regulation were identified later. In humans and other vertebrates, imperfect genome alignments have hindered the discovery of such structures. Here we present RLMHelix, which selects candidate conserved RNA structures from single sequences before any alignment by scoring helices supported by language-model dependencies and thermodynamics. Through distillation, RLMHelix rapidly scans the transcribed human genome and ranks known RNAs higher than conventional thermodynamic scanning. Two-thirds of 15,821 candidates are new to four earlier genome-wide screens, yet agree with in-cell probing, and over half retain structural signal in homologues. In eCLIP data and in exploratory pull-downs, higher RLMHelix scores were associated with more binding by double-stranded RNA-binding proteins. By linking candidates to their binding proteins, RLMHelix yields testable functional hypotheses and illuminates a path from unrecognized human RNA structures to RNA biology and therapeutic targets.

## Introduction

The discovery of new functional RNAs has revealed how RNA controls gene expression and has widened the scope of RNA biology. For RNAs that act through their structure, compensatory substitutions that preserve base pairs are clues to evolutionary constraint.^1^ Comparative genomics, which exploits such patterns of conservation and variation, has led to the discovery of many structured functional RNAs in bacteria.^2–5^ RNA structures are also emerging as therapeutic targets, and small molecules have been reported that bind the FMN riboswitch, the miR-96 precursor and the triple helix at the 3′ end of *MALAT1*.^6–8^ The search for conserved RNA structures (CRSs) thus supports both the discovery of new regulatory mechanisms and RNA-targeted therapeutics.

Searches for CRSs have been highly successful, mainly in bacteria. In humans and other vertebrates, comparative genome screens have also reported many candidate conserved structures.^9^ The main screens used RNAz (thermodynamic stability and structural conservation of aligned sequences), EvoFold (a phylogenetic stochastic context-free grammar, phylo-SCFG, that tests whether substitutions preserve base pairs) and CMfinder (joint inference of an alignment, a structure and a covariance model).^10–12^ Each of these screens predicted tens of thousands to about half a million regions,^13–15^ yet selecting from these predictions the RNAs that have evidence of structural conservation and can be taken forward to functional studies remains difficult.^16^ One reason is that almost all screens depend on genome alignments built on sequence similarity. The substitutions that mark a conserved structure also hinder its alignment. Compensatory substitutions that preserve base pairs lower sequence similarity, so structured RNAs are especially likely to be misaligned or left unaligned.^17–19^ The RNAz and EvoFold screens were therefore confined to regions where alignments are reliable, those that PhastCons marks as highly conserved in sequence (about 4–5% of the human genome).^11,13^ CMfinder infers structures and alignments from unaligned sequences but is computationally demanding.^12^ Even with correct alignments, structural conservation cannot be confirmed by covariation when substitutions are few.^20^

Whereas comparative genomics has found many new classes of functional RNAs in bacteria,^4^ the human CPEB3 and hovlinc ribozymes were discovered by in vitro selection from genomic libraries.^21,22^ This was the case even for the CPEB3 ribozyme, which is conserved across mammals.^23^ More sensitive computational searches might have found such structures from the genome sequence alone, without a selection experiment. What is needed is a computational search that indicates, without relying on alignments, which parts of the genome to examine before any function is measured. ScanFold, which needs no alignment and has been applied across human transcripts,^24^ compares local thermodynamic stability with a shuffled background,^25^ but the structures of functional RNAs cannot always be distinguished from background by stability alone.^26^

Nucleotide dependencies learned by RNA language models offer a way to read conserved base pairs from single sequences.^27–29^ Dependency mapping, which measures how predictions at other positions change when a base is substituted, found previously unreported RNA structures in *Escherichia coli*, supported by in-cell DMS probing and covariation.^27^ If dependencies mark conserved base pairs, they could select candidate conserved structures from single sequences, before any alignment. A dependency map, however, requires 3L + 1 inferences for a sequence of length L, so a scan of the transcribed human genome would take an estimated 680 GPU-days (NVIDIA GB10). Moreover, the human genome is about 700 times longer than that of *E. coli* and its structured RNAs are sparse, so false positives accumulate even for highly specific methods: estimates of the fraction of false predictions in screens with RNAz and EvoFold ranged from 19% to 71% and were higher with backgrounds that preserve dinucleotide composition.^13,30,31^ Base composition in the human genome also varies widely. Together, these constraints require a search of the human genome with dependencies to be fast, to rank candidates against a composition-matched background and to estimate how many of its top candidates would be expected from the background alone.

Here we reverse the order of comparative genomics with RLMHelix, which selects candidate conserved RNA structures from single sequences before any alignment and then tests their conservation in homologues. RLMHelix scores the helices (runs of consecutive base pairs) supported jointly by RNA language-model dependencies and thermodynamic base-pairing probabilities. To make this practical for genome-scale screening, we distilled the dependency map into a model that outputs it in one inference and calibrated the score for base composition. After benchmarking on known RNAs, we scanned the transcribed human genome, compiled the candidates and their vertebrate multiple sequence alignments (MSAs) into the RLM-Fam catalogue, and examined them with evidence independent of the scan: in-cell reactivity, covariation and binding by RNA-binding proteins. We also tested the score in exploratory pull-downs of independently chosen RNAs. RLMHelix ranked known functional RNAs higher than conventional thermodynamic scanning did. Two-thirds of the 15,821 candidates lay in regions that earlier genome-wide screens had not predicted, yet their structures agreed with in-cell probing and were retained in homologues, and the proteins bound to some of them suggest functions to be tested. Together, these results make RLMHelix a scalable way to prioritize unrecognized human RNA structures for functional study.

## Results

### RLMHelix distils dependency maps to propose structures from single sequences

Comparative genomics aligns homologous sequences before looking for structures, so it cannot examine structures in regions that do not align. RLMHelix reverses this order: it proposes structures in single sequences, then collects homologues of the candidates and tests them with evolutionary evidence (Fig. 1a–c). The proposals come from dependency maps, introduced by Tomaz da Silva and colleagues, which measure how much the language model’s prediction at one position changes when the base at another position is substituted;^27^ we use them after average product correction (APC; Methods).^32^ Trained to recover masked nucleotides, RNA language models learn base-pairing constraints without structural labels,^27,28^ so substituting one base of a pair alters the prediction of its partner, and the dependency map reveals base pairs in a single sequence.

**Fig. 1:**
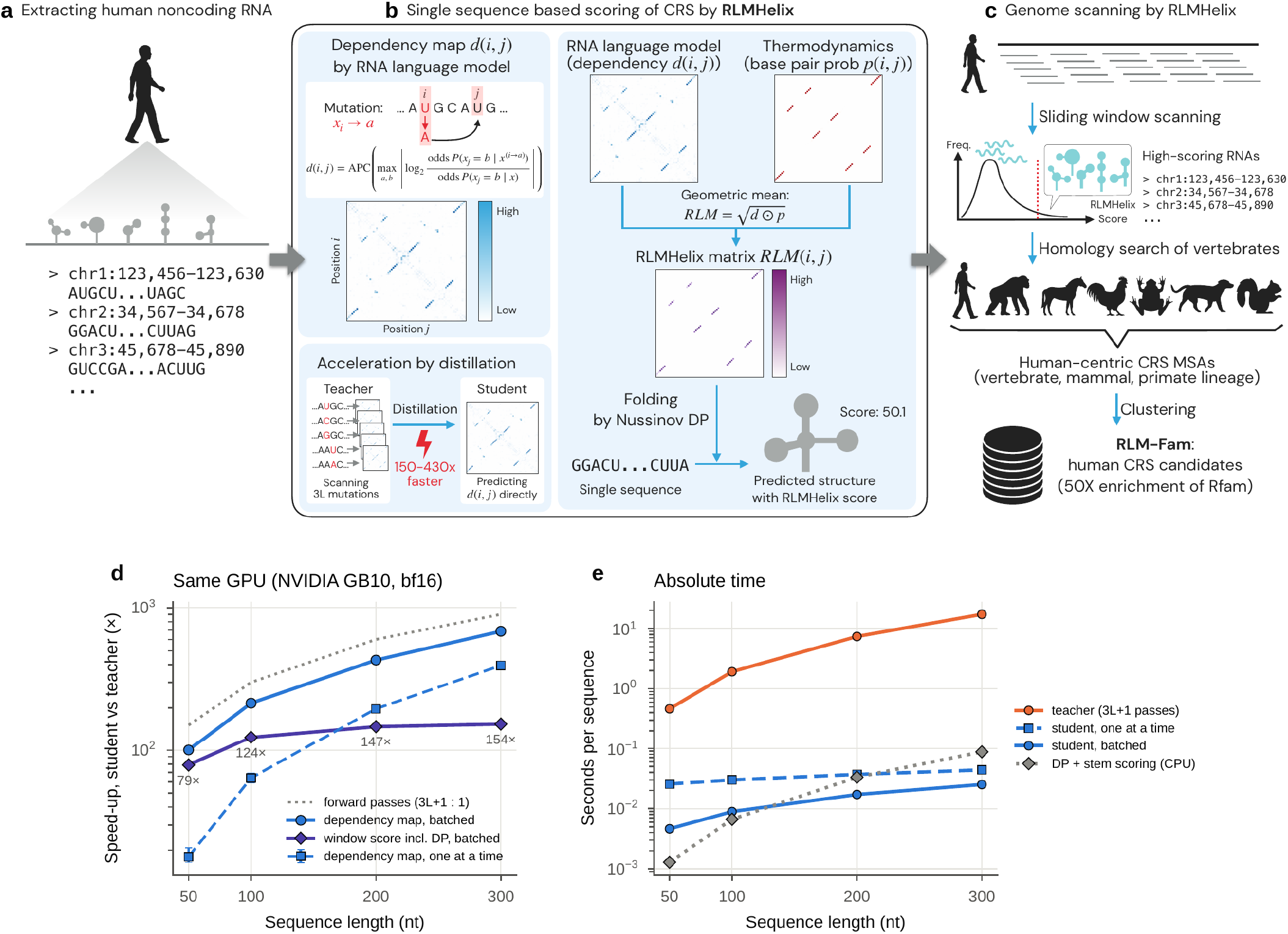
Reversing the order of comparative genomics and distilling dependency maps for fast scanning. **a–c**, Concept (drawn by the authors). **a**, Sequences from the transcribed non-coding regions of the human genome. **b**, Single-sequence scoring by RLMHelix. Substituting one nucleotide at a time in an RNA language model and recording the change in its predictions at other positions gives the APC-corrected dependency map (top left).^27^ Because this map (the teacher) needs 3L + 1 inferences for a sequence of length L, it was distilled into a student that outputs the map in a single inference (bottom left). The geometric mean of the dependency and the RNAfold base-pair probability weights each position pair, non-crossing DP chooses base pairs, and stems of at least four consecutive base pairs are summed into the RLMHelix score (right). **c**, Genome scanning: windows are ranked by the composition-calibrated score, homologues of the top windows are aligned across vertebrates, and the candidates are grouped into RLM-Fam. **d**, Speed-up of the student over the teacher on the same GPU, for 20 windows each of 50, 100, 200 and 300 nt from transcribed regions of chr22. Dependency maps were computed one at a time (median and interquartile range) or 64 at a time (batched); purple line and numbers, speed-up of whole-window scoring including the DP (batched); dotted line, ratio of inference counts. **e**, Time per sequence. At 200 nt, the teacher takes 7.4 s and the student 0.037 s one at a time (0.017 s batched); the DP and stem scoring on the CPU take 0.033 s.

The RLMHelix score summarizes the support for helices from dependencies and the thermodynamic basepair probability (BPP; Fig. 1b). Each position pair is weighted by the geometric mean of its dependency and BPP, Nussinov-type dynamic programming (DP) chooses the non-crossing base pairs with the largest total weight,^33^ and the weights of stems of at least four consecutive pairs are summed. The score is a heuristic rather than a probabilistic model of folding; it is highest for long helices supported by both the language model and thermodynamics.

Distillation made dependency maps feasible genome-wide. The original dependency mapping (the teacher) needs 3L + 1 inferences for a sequence of length L,^27^ so we trained a small model (the student) that outputs the APC-corrected map in a single inference from the representations of the frozen language model (knowledge distillation;^34^ Fig. 1b). For 200-nucleotide (nt) windows, with the student run in batches of 64 on the same GPU as the teacher, scoring including DP was 147 times faster than with the teacher (Fig. 1d, e), shortening a scan of about 7.9 million windows of the transcribed human genome from an estimated 680 GPU-days to about 2 days on a single desktop GPU (NVIDIA GB10). On 5,313 bpRNA-new sequences, with each map combined with the BPP in the same way, the student predicted base pairs with an F1 of 0.61, against 0.56 for the teacher and 0.62 for the RNAfold MFE structure (Supplementary Fig. S1).^35^

### Dependencies add discriminative information beyond thermodynamics

Thermodynamic stability is the standard clue in searches for structured RNAs, but it separates functional RNAs poorly from background.^26^ To test whether the RLMHelix score adds information to the BPP, we compared the human known RNA of each of 328 vertebrate Rfam families with 100 dinucleotide-preserving shuffles of that RNA, using precision–recall curves because each family has a single positive (Fig. 2a, Methods, Supplementary Table S1).

**Fig. 2:**
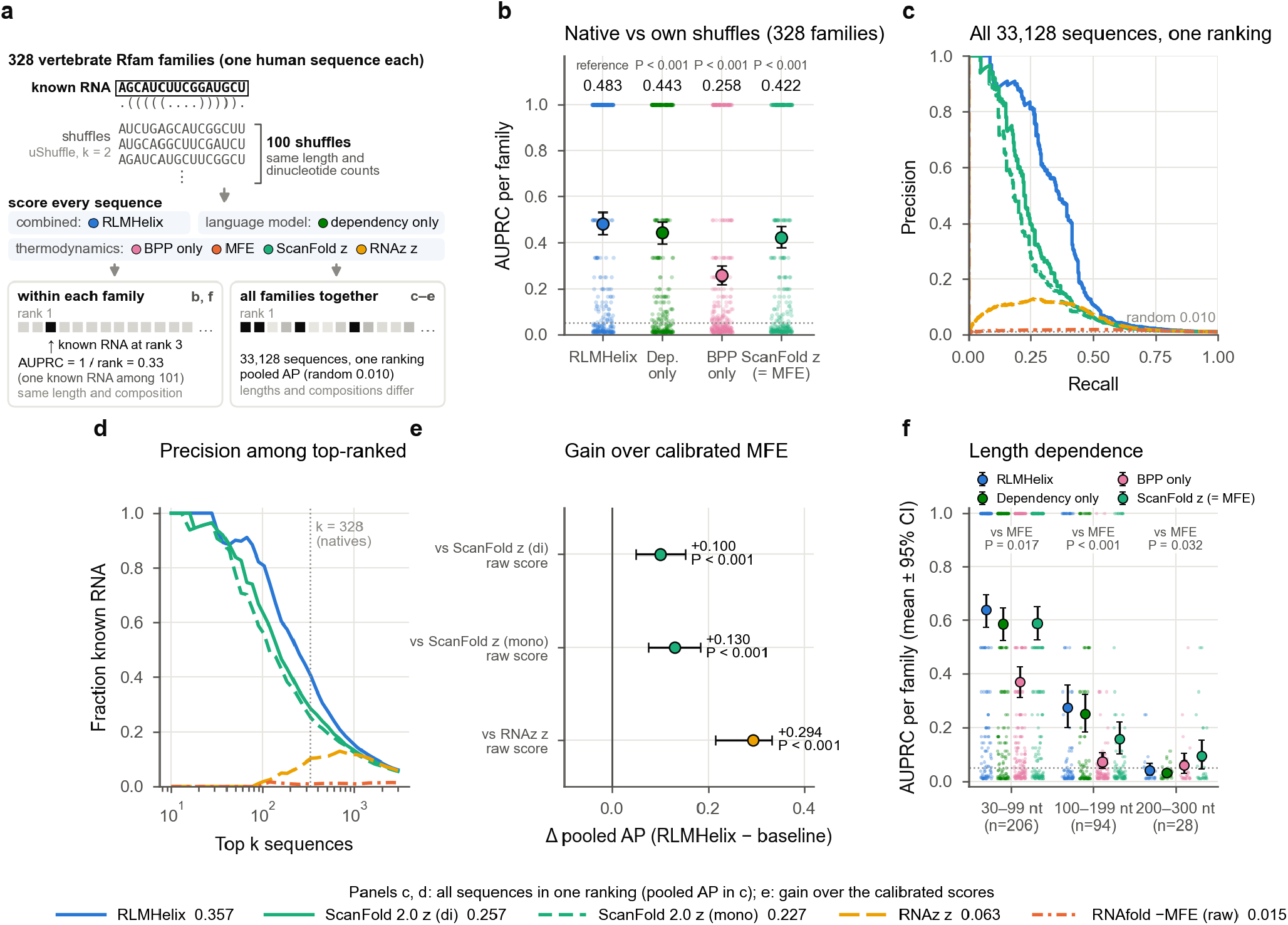
Dependencies add discriminative information beyond thermodynamics. For 328 vertebrate Rfam families, the known human RNA of each family was compared with 100 shuffles of the same dinucleotide composition (uShuffle, k = 2).^36^ **a**, Design of the benchmark (schematic; the sequences are illustrative). **b**, AUPRC of each family’s known RNA against its own shuffles (1/rank, with one positive per family). Small points, families; circles and bars, mean and 95% interval; numbers, means, with the paired P against RLMHelix above; dotted line, chance expectation (0.051). Sequence length and composition are identical within a family, so the ScanFold 2.0 z-score and the raw MFE rank the sequences identically. Difference RLMHelix − MFE, +0.061 (95% interval 0.027–0.094). **c**, Precision–recall curves for all 33,128 sequences in a single ranking; the key below the figure gives the pooled AP. ScanFold 2.0 z-scores^25^ and RNAz z-scores are relative to the MFE distribution of shuffles predicted from composition (Supplementary Methods); the raw RNAfold −MFE, which depends strongly on length, is drawn for comparison. **d**, Fraction of known RNAs among the top k sequences (dotted line, k = 328, the number of known RNAs). **e**, Differences in pooled AP between the raw RLMHelix score and calibrated thermodynamic scores, with the paired P beside each. The 95% intervals are 0.048–0.153 against ScanFold 2.0 z (dinucleotide), 0.075–0.183 against the same (mononucleotide) and 0.214–0.333 against RNAz z. **f**, AUPRC by sequence length, as in **b**, with the P of RLMHelix against the MFE in each length range. Means for RLMHelix and the MFE: 0.638 and 0.588 (30–99 nt), 0.275 and 0.158 (100–199 nt), 0.042 and 0.093 (200–300 nt). Intervals are 95% bootstrap intervals over families (2,000 resamples in **b** and **f**, 1,000 in **e**). P values are from two-sided paired bootstrap tests over families (n = 328 families; 206, 94 and 28 in **f**), computed as twice the fraction of resampled differences on the smaller side of zero (10,000 resamples in **b** and **f**, 2,000 in **e**; P < 0.001, no resample crossed zero), and are not adjusted for multiple comparisons.

Within families, RLMHelix ranked the known functional RNA higher than the RNAfold minimum free energy (MFE) did: the mean area under the precision–recall curve (AUPRC) was 0.48 against 0.42 (Fig. 2b). Because these sequences share length and composition, the composition-calibrated ScanFold 2.0 z-score ranks them as the MFE does.^25^ Weighting pairs by the dependency map alone or by the BPP alone (ablations) gave a lower mean AUPRC than RLMHelix (Fig. 2b), so dependencies and thermodynamics carry complementary information.

We next tested whether the advantage persists when sequences of different lengths and compositions are ranked together, as in a real scan. Among all 33,128 sequences (Fig. 2c–e), raw MFE depended strongly on length, whereas RLMHelix reached an average precision (AP) of 0.36, against 0.26 for the ScanFold 2.0 z-score. Of the top 328 sequences (the number of known RNAs), 134 were known RNAs with RLMHelix and 95 with ScanFold 2.0. The known RNAs among the ScanFold 2.0 top 328 were almost all miRNA precursors, whereas those among the RLMHelix top 328 also included snoRNAs (13 versus 0; Supplementary Fig. S2).

Despite outperforming thermodynamics on short structured RNAs, RLMHelix did not distinguish long RNAs from their shuffles: for the 28 families of 200–300 nt its AUPRC was indistinguishable from chance and below that of the MFE (Fig. 2f), and the 127 Rfam families annotated as lncRNA were barely distinguished by any score (Supplementary Fig. S2). The gain from dependencies thus lies mainly in structured RNAs shorter than 200 nt with well-defined stems.

### The same dependency representation scores multiple sequence alignments

Whether a candidate chosen from a single sequence is a conserved structure has to be tested in an alignment of its homologues. We therefore predicted a dependency map with the student for each sequence of the alignment, averaged the maps for each pair of alignment columns and combined them with the RNAalifold BPP in the same DP (RLMHelix-MSA; Fig. 3a), comparing each alignment with 100 SISSIz shuffles that preserve phylogeny and composition.^37^ Selection on single sequences and testing on alignments then use the same representation.

**Fig. 3:**
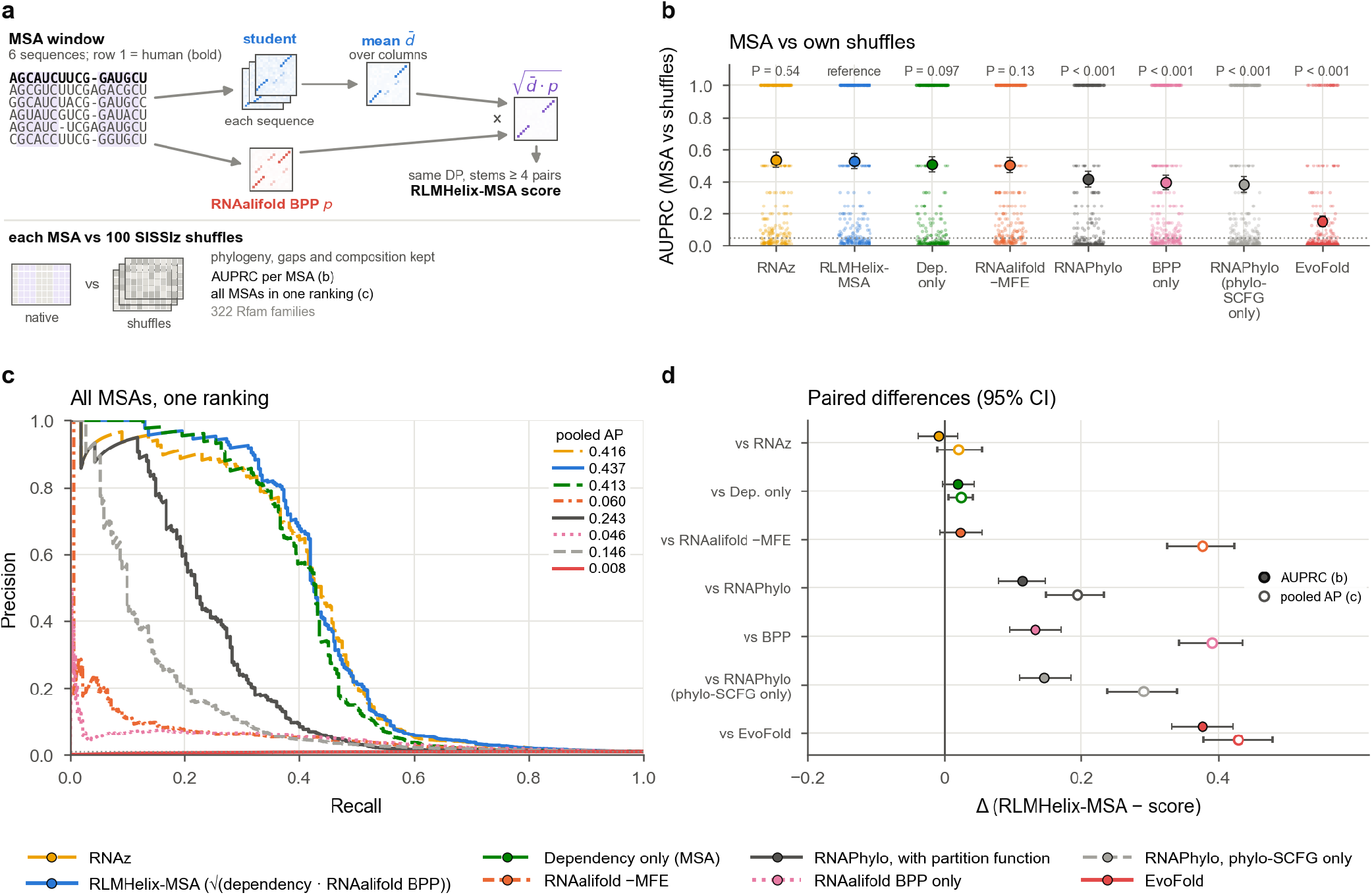
The same dependency representation scores multiple sequence alignments. Rfam, 322 human-centred families (the 328 of Fig. 2 without six miRNA families whose sequences are nearly identical across species, so that the MSA carries no information). Each MSA was compared with 100 SISSIz shuffles that preserve phylogeny and composition.^37^ **a**, RLMHelix-MSA (schematic; the matrices and sequences are illustrative). The student predicts a dependency map for each sequence of the MSA pruned to six sequences (RNAz scores the same six), and the maps are averaged for each pair of alignment columns. The geometric mean of the averaged map and the RNAalifold base-pair probability is scored with the DP and stem summation of Fig. 1b. **b**, AUPRC of each family’s MSA against its own shuffles (points) and the mean with 95% interval, with the scores ordered by the mean (dotted line, chance expectation 0.051). Above, two-sided paired bootstrap P against RLMHelix-MSA (10,000 resamples of families). RNAPhylo weights phylo-SCFG posteriors by the thermodynamic partition function, and RNAPhylo (phylo-SCFG only) omits it.^15^ **c**, Precision–recall curves for all MSAs in a single ranking; the legend in the panel gives the pooled AP of each curve (line styles as in the key below the figure). The raw RNAalifold −MFE depends on length, which lowers its pooled AP. **d**, Paired differences between RLMHelix-MSA and the other scores (filled, mean AUPRC of **b**; open, pooled AP of **c**). Intervals are 95% bootstrap intervals over families (10,000 resamples for mean AUPRC and 2,000 for pooled AP). P values are from two-sided paired bootstrap tests over the 322 families, computed as twice the fraction of resampled differences on the smaller side of zero (P < 0.001, no resample crossed zero), and are not adjusted for multiple comparisons. The values of all scores and their differences are in Supplementary Table S2; comparisons on the predictions of earlier screens are in Supplementary Fig. S3.

On 322 of the 328 vertebrate Rfam families (excluding six miRNA families whose sequences are nearly identical across species), RLMHelix-MSA, a heuristic score not trained for this task, discriminated known RNAs about as well as the RNAalifold free energy (RNAalifold −MFE) and RNAz, a support vector machine trained on alignments of known non-coding RNAs and their shuffles:^10,38^ its mean AUPRC within families (0.53) was comparable to that of RNAz (0.54) (Fig. 3b, d). With all alignments ranked together, its AP of 0.44 was similar to that of RNAz (0.42) and far above that of the BPP-only ablation, a gain that comes from the dependencies (Fig. 3c).

By contrast, the phylogenetic models used by earlier genome screens scored lower on both measures (Fig. 3b– d, Supplementary Table S2). The best of them, RNAPhylo, which weights phylo-SCFG posteriors by the thermodynamic partition function and was used to score the CMfinder screen,^15^ reached a mean AUPRC of 0.41; RNAPhylo with the phylo-SCFG alone and EvoFold scored lower still. Thermodynamics thus improved the phylo-SCFG, as it improved the dependencies (Fig. 3b). EvoFold also tended to score shuffled alignments highly, as reported earlier.^39^ Supplementary Fig. S3 applies the same comparison with shuffles to regions predicted by earlier screens (EvoFam and RNAz 2005), which measures agreement with those screens rather than accuracy. RLMHelix can therefore score not only single sequences but also MSAs built by other comparative-genomics pipelines, discriminating known RNAs about as well as RNAz.

### RLM-Fam construction by genome-wide scanning and two-stage selection

Having found that the score separates known RNAs from shuffles in single sequences and alignments, we scanned the transcribed non-coding regions of the human genome in 200-nt windows with a 100-nt step, scoring 7,869,821 windows after filtering repeats and other regions (Fig. 4a). With the raw score, AT-rich windows accumulated at the top (42% of the top 0.1% had a GC content below 35%, against 29% of random windows; Fig. 4b), so we ranked windows by P values calibrated with a generalized extreme value (GEV) model that predicts the score distribution of dinucleotide-preserving shuffles from sequence length and composition. Calibration lowered the fraction of AT-rich windows in the top 0.1% to 17% while largely preserving the overlap with known RNAs.

**Fig. 4:**
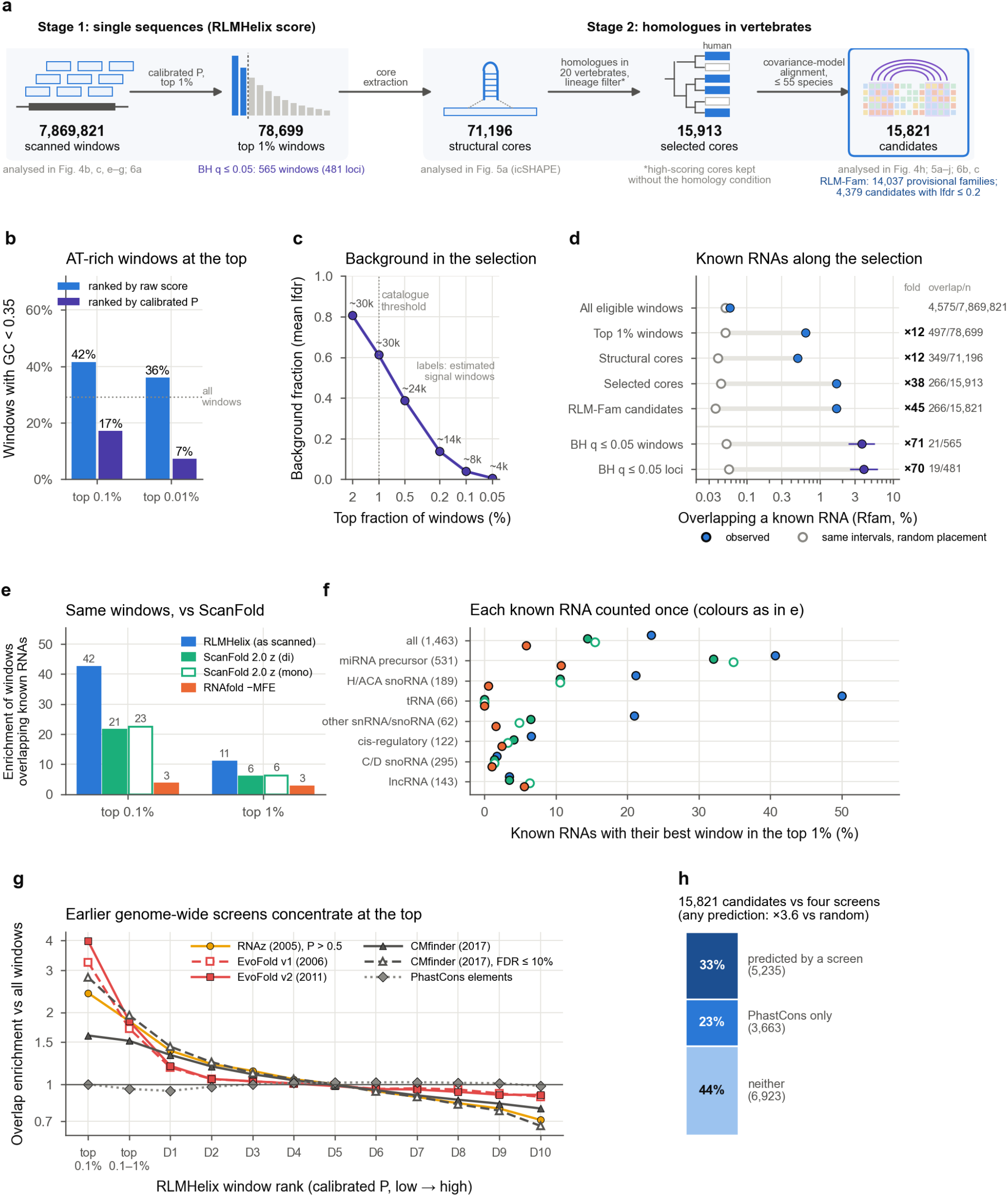
Scanning the genome concentrates known RNAs and earlier predictions and yields candidates beyond them. Windows of 200 nt (100-nt step) of the transcribed non-coding human genome. **a**, Two-stage selection of candidates (schematic; the glyphs are illustrative and the numbers are counts). Filled boxes in the tree, species with homologues. Below each set, its stricter subsets and, in grey, the panels that analyse it. The pull-down baits (Fig. 6d–f) were chosen independently of the scan. **b**, Fraction of windows with a GC content below 35% at the top of the ranking (n = 7,870 and 787 windows); dotted line, 200,000 random windows. **c**, Fraction of background windows (mean lfdr) by the top fraction of windows under Efron’s empirical null^40^ (Supplementary Fig. S4); labels, estimated numbers of signal windows. Values beyond the top 0.1% extrapolate the null into the tail. **d**, Fraction of intervals overlapping a known RNA (Rfam 15.1, same strand)^41^ at each stage of selection, with Wilson 95% intervals (at right, the fold over random placement and overlap/n, with n the intervals of each set); open circles, intervals of the same lengths placed at random in the scanned space (mean of 20–1,000 placements); violet, BH q ≤ 0.05. **e**, Enrichment of windows overlapping known RNAs among the top windows (n = 7,870 and 78,699 windows; 4,575 of all windows overlap a known RNA), for the calibrated RLMHelix score used for the scan and for thermodynamic scores computed on the same windows. **f**, Each of 1,463 known RNAs (Rfam hits at least half covered by windows) counted once: fraction with its best window in the top 1%, by RNA class (numbers in parentheses; ‘all’ also includes RNAs of classes not shown separately). Overall, 23.3% for RLMHelix and 14.4% for ScanFold 2.0 (dinucleotide); difference +8.9 percentage points (95% interval 6.5–12.0; bootstrap over 1-Mb blocks, 200 replicates). **g**, Fold enrichment, by window rank, of the overlap (either strand) with four earlier screens: RNAz 2005 (Set1, P > 0.5),^13^ EvoFold (v1 and v2)^11,14^ and the CMfinder CRSs (all with RNAPhylo score > 50, and those with a reported FDR ≤ 10%).^15^ D1–D10, deciles of the windows below the top 1% (D1 highest). PhastCons conserved elements (hg17),^48^ which approximate the input of RNAz 2005 and EvoFold, serve as a control. **h**, The 15,821 RLM-Fam candidates by their overlap with the four prediction sets (either strand); ‘PhastCons only’, no prediction but within PhastCons conserved elements; heading, fold enrichment of the overlap with any of the four sets over intervals of the same lengths placed at random in the scanned space (Supplementary Fig. S7a).

The scan score is a single-sequence signal, so candidates were selected in two stages: the top 1% of windows by the calibrated P value, then homology (Fig. 1c and Fig. 4a). The structural cores of these windows (the outermost base-paired intervals of their predicted structures; 71,196 after overlapping windows were merged and identical cores collapsed; Methods) were searched for homologues in 20 vertebrates and filtered with lineage-based conditions (15,913; cores with especially high scores were kept without the homology condition). After the search was extended to 34 further vertebrates, the 15,821 candidates that could be aligned with covariance models (up to 55 species, including human; Methods) were grouped by reciprocal searches into 14,037 provisional families, which form RLM-Fam (Fig. 4a, d); we refer to candidates by family identifiers such as RLMFam00092.

We then estimated how many of the top windows carry signal. An empirical null fitted to the centre of the distribution of z-transformed P values (Efron’s method)^40^ attributed about 30,000 of the top 1% of windows (78,699) to signal (Fig. 4c, Supplementary Fig. S4). From this null we computed the local false discovery rate (lfdr) of each window and took lfdr ≤ 0.2 (an estimated probability of at most 20% that the window belongs to the background) as statistical support (4,379 RLM-Fam candidates). Because these estimates extrapolate the null into the tail, the 565 windows (481 loci; Methods) with a nominal Benjamini–Hochberg (BH) q ≤ 0.05 serve as a stricter reference set. RLM-Fam was not selected at a fixed false discovery rate; the lfdr is used to prioritize candidates, and the false discovery rate of RLM-Fam is estimated from homologues in the next section.

If the selection enriches for structured RNAs, overlap with known RNAs should increase at each stage. Same-strand overlap with known RNAs (Rfam 15.1)^41^ rose to 12, 45 and 70 times that of randomly placed intervals of the same lengths for the top 1% of windows, RLM-Fam candidates and the 481 BH loci (Fig. 4d, Supplementary Fig. S5). RNAs with several stable stems or long helices, such as tRNAs, miRNA precursors and H/ACA snoRNAs, were enriched more strongly than short cis-regulatory hairpins and C/D snoRNAs, as expected for a score that sums stems (Supplementary Fig. S6). Known functional cis-acting structures were enriched about fourfold in the top 1%, and five of the 67 in the scanned space entered RLM-Fam, against 0.8 expected (Supplementary Table S3): the METTL16-target hairpin hp1 of *MAT2A*,^42^ the iron-responsive elements of *TFRC* and *EPAS1*,^43,44^ the triple helix of *MALAT1*^45,46^ and a histone mRNA stem-loop.^47^

The advantage over thermodynamic scores observed on Rfam also held in the scan itself. Because the Rfam comparison used shuffles rather than the windows of a real scan, we scored the same windows with RNAfold MFE and ScanFold 2.0 z-scores (Methods). In the top 0.1%, the calibrated RLMHelix enriched windows overlapping known RNAs 42-fold, against 21-fold for ScanFold 2.0 and less for raw MFE (Fig. 4e). Of the known RNAs, 23.3% had their best window in the top 1%, against 14.4% with ScanFold 2.0; the gain came mostly from miRNA precursors, tRNAs and H/ACA snoRNAs, and half of the tRNAs reached the top 1% with RLMHelix but none with ScanFold 2.0 (Fig. 4f).

We compared the candidates with the predictions of four genome-wide screens (the RNAz screen,^13^ two versions of EvoFold^11,14^ and the CMfinder screen for conserved structures^15^). Candidates overlapped the predictions of each screen three- to tenfold more often than intervals of the same lengths placed at random, and 5,235 (33%) overlapped at least one prediction. Yet two-thirds of the candidates (10,586 of 15,821) lay where no earlier screen had predicted a structure, 3,336 of these with statistical support, and 6,923 of these 10,586 (44% of all candidates) also lay outside PhastCons conserved elements, which approximate the regions that the 2005 RNAz screen and EvoFold examined (Fig. 4g, h, Supplementary Fig. S7). The scan thus overlaps many earlier predictions and proposes many candidate structures where alignment-based screens predicted none.

### Candidate structures are supported by in-cell reactivity and in homologues and, for some, by covariation

Enrichment for known RNAs raises the question of whether the structures of the 98% of candidates outside Rfam are also supported. We tested them with three kinds of evidence independent of the scan: in-cell reactivity, structural signal in homologues and covariation. If the predicted structures form in cells, bases predicted to be unpaired should be more reactive than paired ones; we measured this as the area under the receiver operating characteristic curve (AUROC) separating unpaired from paired nucleotides by reactivity. In public icSHAPE data (GSE145805, five cell lines),^49^ this held for the 757 measurable cores from the top 1% of windows (474 of them RLM-Fam candidates; mean AUROC 0.67, against 0.50 for controls with the structure shifted along the sequence; Fig. 5a). The agreement was slightly lower than with RNAfold MFE structures (0.68) and the same as with structures selected with BPP only. Agreement per candidate in GSE145805 correlated with that in a second icSHAPE data set (GSE74353; Spearman ρ = 0.76; Fig. 5b).^50^ For candidates that overlapped no earlier prediction and lay outside PhastCons conserved elements, agreement was slightly weaker but similar (mean AUROC 0.64; Fig. 5a). Adding dependencies thus kept the predicted structures about as consistent with in-cell structure as thermodynamic predictions.

**Fig. 5:**
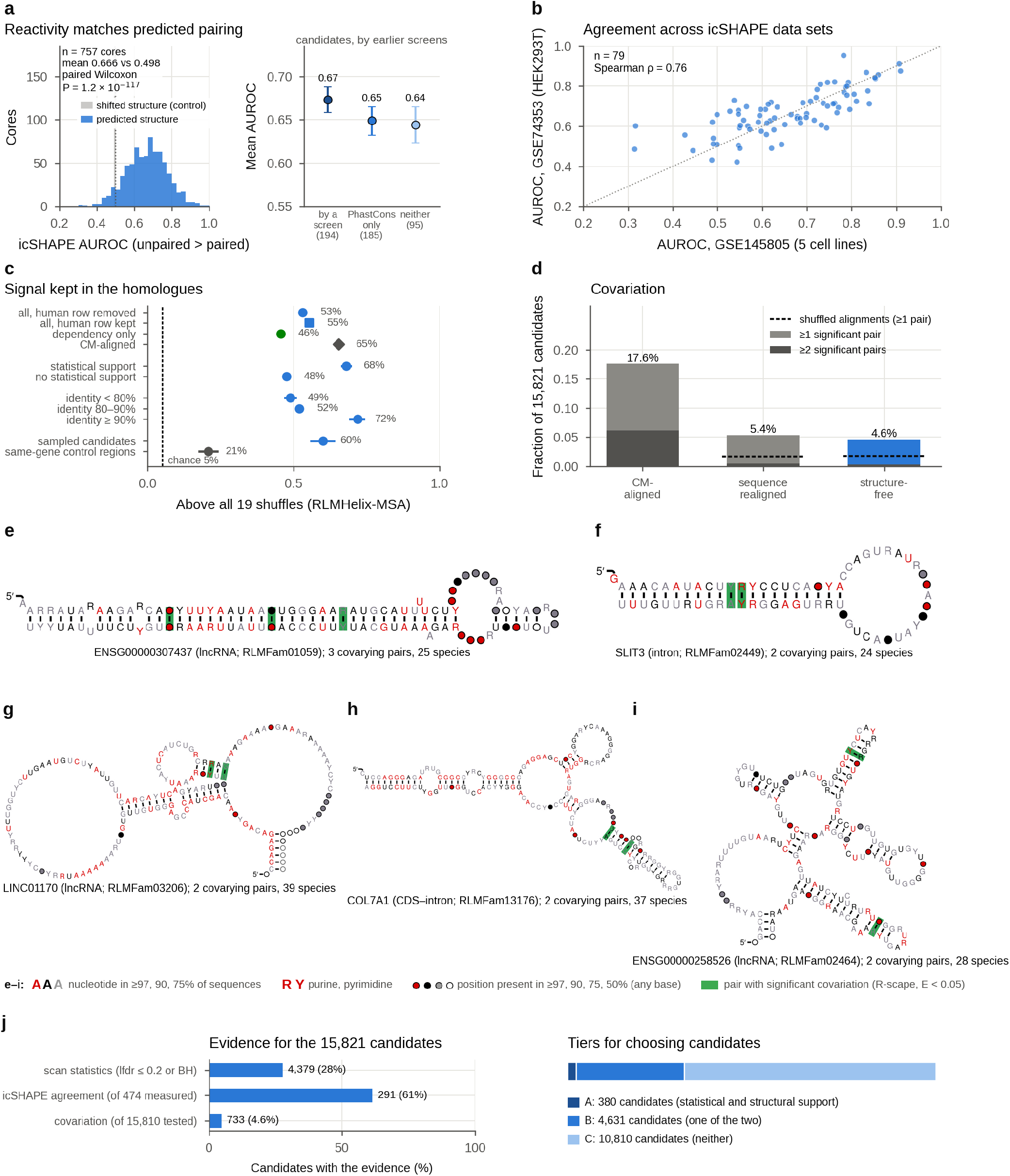
Candidate structures are supported by in-cell reactivity and in homologues and, for some, by covariation. **a**, icSHAPE agreement of the 757 measured cores (474 of them RLM-Fam candidates; AUROC averaged over the cell lines of each core): AUROC for higher reactivity at nucleotides predicted to be unpaired (GSE145805, five cell lines),^49^ against a control with the structure shifted along the sequence (grey); two-sided paired Wilcoxon signed-rank test over cores. Structures from the RNAfold MFE give a mean of 0.683, and structures chosen with BPP only 0.666. Right, the measured RLM-Fam candidates by their relation to the four earlier screens (groups as in Fig. 4h, n in parentheses; mean and 95% interval from 2,000 bootstrap resamples of candidates). **b**, Agreement per candidate between GSE145805 and GSE74353 (HEK293T; n = 79 candidates; Spearman’s ρ, two-sided P = 2 × 10^−16^).^50^ **c**, Fraction of candidates whose alignment of homologues scores above all 19 SISSIz shuffles with RLMHelix-MSA (95% intervals from 5,000 bootstrap resamples of candidates; dashed line, 5% by chance). In the main condition (top row; 14,565 candidates, as in the next two rows, which keep the human row or score with the dependencies alone, without the RNAalifold BPP), homologues found without using the structure were aligned with MAFFT^52^ and the human sequence was removed. Covariance-model alignments (14,356 candidates) follow the predicted structure, which can inflate the signal. Below, by statistical support in the scan (lfdr ≤ 0.2; 3,981 and 10,584 candidates) and by alignment identity (3,329, 9,907 and 1,320), and, at the bottom, 600 randomly chosen candidates and a low-score region in the same gene of each, searched, aligned and scored in the same way (593 pairs). **d**, Fraction of the 15,821 candidates with ≥1 or ≥2 predicted base pairs showing significant covariation (R-scape, E < 0.05)^51^ under three alignment conditions; dashed lines, the same call on one SISSIz shuffle of each realigned and structure-free alignment (1.7%; no shuffles were made for the covariance-model alignments). Blue, the structure-free condition used in the text. **e–i**, R2R consensus diagrams^53^ of RLM-Fam candidates with significantly covarying base pairs in structure-free alignments, chosen among those that do not overlap Rfam and have low copy numbers (symbols as in the key below). **j**, Evidence for the 15,821 candidates (left; icSHAPE agreement among the 474 measured candidates, covariation among the 15,810 tested) and the tiers built from it (right): A, statistical support (lfdr ≤ 0.2 or a BH window) and structural support (icSHAPE or covariation in structure-free alignments); B, one of the two; C, neither (380, 4,631 and 10,810 candidates; structural support for 1,012; Supplementary Table S4).

We next asked whether the structures are conserved in evolution, which in-cell reactivity cannot show. With RLMHelix-MSA, we compared structure-free alignments (homologues retrieved without the structure and aligned with MAFFT, with the human sequence removed) with 19 SISSIz shuffles that preserve phylogeny, gap pattern and dinucleotide content.^37^ The alignment scored above all 19 shuffles for 53% of 14,565 candidates, against 5% expected by chance (Fig. 5c); alignments built with a covariance model of the predicted structure gave 65%, but alignment to the structure can inflate the signal. From the rank of each alignment among its 19 shuffles (Storey’s method; Methods), 13% of candidates are expected to have alignments indistinguishable from their shuffles, which we take as the false discovery rate of RLM-Fam against shuffled alignments, the background used by earlier genome-wide screens. Shuffles do not capture everything that real alignments share, however: low-score regions in the same genes as 600 randomly chosen candidates scored above all 19 shuffles in 21% of cases, against 60% for those candidates (Fig. 5c), and with these regions as the background the estimate rises to 25%. The false discovery rate of RLM-Fam thus lies between about 13% and 25%, depending on the background (Supplementary Methods). The fraction above all shuffles was 68% for candidates with statistical support and 48% for the others, and 49% even for alignments below 80% identity. More than half of the candidates thus keep their structural signal in non-human homologues, but homologues also share sequence with the human sequence, so this signal alone does not separate constraint on structure from sequence conservation; covariation, which requires compensating substitutions, is the stricter evidence.

In structure-free alignments, 4.6% of candidates (733 of the 15,810 tested) had at least one predicted base pair with significant covariation (R-scape),^51^ about three times the rate in SISSIz shuffles of the same alignments (4.7% versus 1.7% among the 14,577 candidates whose alignments could be shuffled; Fig. 5d); alignments built with a covariance model of the predicted structure (CM-aligned) gave 17.6%, and the same sequences realigned by sequence alone (sequence-realigned) 5.4%. A base pair that has not varied carries no covariation by definition, so the absence of covariation does not show that a structure is absent,^1,20^ but the candidates contain structures supported by covariation at a rate that shuffles do not produce (examples in Fig. 5e–i). Covariation was also detected about twice as often among candidates with low lfdr (7.3% versus 3.6%; Supplementary Fig. S8), so the scan statistics agree with independent evidence; Fig. 5j and Supplementary Table S4 summarize the evidence for each candidate.

### Double-stranded RNA-binding proteins bind RLM-Fam candidates, and individual candidates yield testable hypotheses

Having compiled the candidates and their evidence, we next sought to link them to function through their binding proteins, because many structured RNAs act through RNA-binding proteins (RBPs). We matched candidates with the eCLIP binding sites of 168 RBPs measured by ENCODE in K562 and HepG2 cells.^54^ With all scanned windows as background, as in an earlier analysis of conserved structures,^15^ 42% of the 250 experiments were enriched on candidates, but much of this enrichment reflected genomic location: against low-score regions in the same gene, strand and kind of region, matched for exon overlap and GC content, enrichment remained in 12% (30 experiments; Supplementary Fig. S9, Supplementary Table S5).

We then examined which proteins were most strongly enriched after matching. Double-stranded RNA-binding proteins (dsRBPs) together were enriched on candidates (odds ratio 2.2; Supplementary Fig. S9b), led by DGCR8, the RNA-binding component of the Microprocessor; much of this came from candidates near known miRNA genes, without which the odds ratio was 1.5 (P = 0.06). Across windows stratified by gene, region type, exon overlap and GC content (Methods), dsRBP binding increased with the score (odds ratio 8.9 for windows with calibrated P < 0.001 relative to those with calibrated P > 0.5), even when windows near miRNA genes were excluded, whereas binding by the other 162 RBPs decreased (Fig. 6a). The binding that rises with the score is concentrated in proteins that read duplexes, consistent with the long helices of many candidates.

**Fig. 6:**
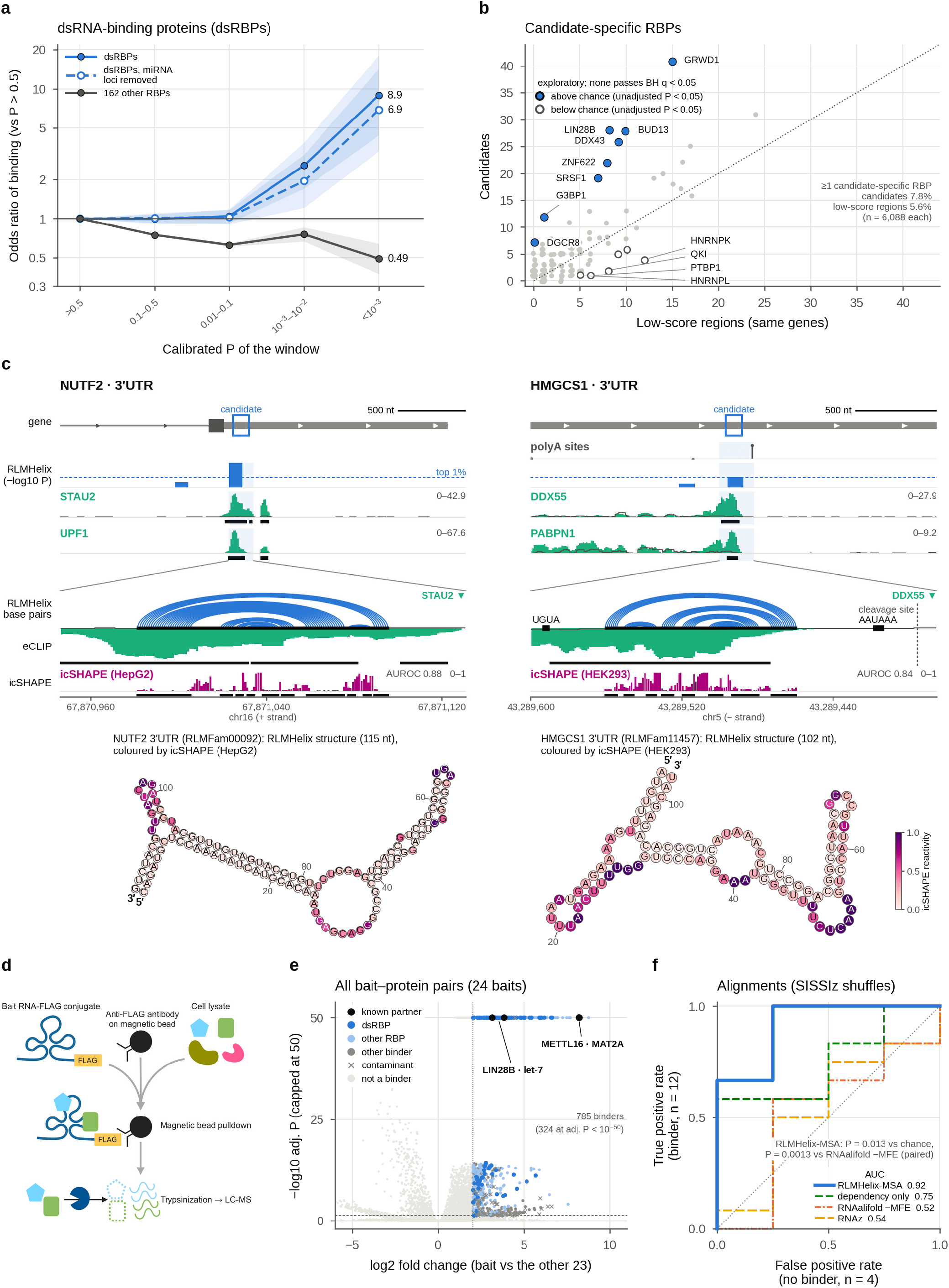
Protein binding links candidates to testable hypotheses, and in pull-downs of independently chosen RNAs, higher scores accompany structure-recognizing binders. **a**–**c**, ENCODE eCLIP (250 experiments, 168 RBPs, K562 and HepG2).^54^ **a**, Mantel–Haenszel odds ratios (95% intervals) for binding by six double-stranded RNA-binding proteins (DGCR8, DROSHA, ILF3, STAU2, DHX30 and XPO5) and by the other 162 RBPs, by the calibrated P value of the window, relative to windows with P > 0.5 (n = 690,368 windows; 3,238,180, 537,727, 17,394 and 5,269 in the bins of P 0.1–0.5, 0.01–0.1, 0.001–0.01 and < 0.001; strata in Methods). Dashed line, the same six proteins after removing windows within 500 nt of a GENCODE miRNA gene. **b**, Candidate-specific RBPs (binding the candidate and none of ≥2 same-gene controls, with the strongest peak in the gene at the candidate), counted at candidates and at low-score regions of the same genes. Each point is an RBP; tests are two-sided exact McNemar tests on 6,088 pairs of a candidate and a low-score region, with BH adjustment across RBPs. **c**, Two unannotated 3′UTR candidates, *NUTF2* (STAU2 and UPF1; PCBP1 not drawn) and *HMGCS1* (DDX55 and PABPN1). Each column runs 5′→3′ from left to right (coordinates reversed for *HMGCS1*, on the minus strand). Above, HepG2 eCLIP read coverage, not crosslink sites (mean of two replicates; grey line, input; black bars, reproducible (IDR) peaks). Gene model, MANE Select (blue box, core; arrows, direction of transcription); polyA sites, PolyASite 2.0 cleavage sites (height, share among the sites in view);^57^ RLMHelix, windows in the top 5% (dashed line, top 1%). The zoom shows the base pairs of the RLMHelix structure, the signal of the first RBP and icSHAPE reactivity (the source study’s icSHAPE-pipe scores; HEK293 for *HMGCS1*, which was not measured in HepG2; black bars, nucleotides paired in the RLMHelix structure). Its AUROC (unpaired versus paired nucleotides) is for the cell line drawn; the text gives the mean over cell lines. Below, the same RLMHelix structures coloured by the same reactivity. **d**, Pull-down mass spectrometry (schematic; shapes are illustrative). **e**, All bait–protein pairs of the 24 baits (exploratory). Each point is one bait–protein pair, with the log2 fold change and P against the other 23 baits from a proDA Wald test (measurements in Supplementary Methods; P adjusted by Benjamini–Hochberg across proteins for each bait; −log10 capped at 50); pairs beyond both lines (adjusted P = 0.05, dashed; log2 fold change = 2, dotted) are binders. LIN28B was recovered on both let-7 baits and METTL16 on both *MAT2A* baits; the known partners of SNORA2, the miR-9-2 precursor and the histone stem-loop were not binders. Plots for each bait are in Supplementary Fig. S12. **f**, ROC curves of the shuffle z of each alignment score (SISSIz shuffles) for the 16 human baits of ≥30 nt, separating the 12 with a structure-recognizing binder (a double-stranded RNA-binding protein or a known partner that reads a sequence in a hairpin loop) from the 4 without. P values in the panel: two-sided Mann–Whitney test against chance (n = 12 and 4 baits) and two-sided paired DeLong test against RNAalifold −MFE, not adjusted for multiple comparisons. The 95% interval of the AUC of RLMHelix-MSA from 10,000 bootstrap resamples of baits is 0.68–1.00. Single-sequence scores are in Supplementary Fig. S13c.

Beyond enrichment across candidates, we sought RBPs that bind individual candidates specifically, but eCLIP binding alone did not identify such partners. Candidate-specific RBPs, which bind none of the same-gene controls and whose strongest peak in the gene lies on the candidate (Fig. 6b), were found for 7.8% of candidates but also for 5.6% of low-score regions in the same genes; this chance ratio (0.72) implies that about 70% of partners assigned from eCLIP alone could arise by chance (chance ratio 0.45 for dsRBPs; Supplementary Fig. S10). Restricting calls to those consistent with a known property of the partner did not separate specific binding from chance either, because features that these properties require, such as long helices, differ in frequency between candidates and controls (Supplementary Fig. S10). The partner of an individual candidate therefore cannot be assigned from these statistics and has to be argued case by case, combining binding with the partner’s properties and known examples.

Two unannotated structures in 3′ untranslated regions (UTRs) show how the catalogue leads to hypotheses about individual functions (Fig. 6c). In the *NUTF2* 3′UTR (RLMFam00092; an 18-base-pair helix; icSHAPE agreement 0.86; outside the predictions of the four earlier screens), strong Staufen2 (STAU2) and UPF1 signals lay over the 5′ part of the structure, these two proteins and PCBP1 bound none of the same-gene controls, and nucleotide reactivity followed the predicted pairing (Supplementary Fig. S11f, g). Staufen1 binds duplexes in 3′UTRs and recruits UPF1 to degrade the mRNA (Staufen-mediated mRNA decay, SMD),^55^ and Staufen2 takes part in the same pathway.^56^ With the duplex reader and the decay factor on the same helix, we hypothesize that this structure makes *NUTF2* mRNA a target of SMD; a contribution of its many paralogues and pseudogenes to these binding and reactivity signals remains to be excluded. In the *HMGCS1* 3′UTR (RLMFam11457; 102 nt; icSHAPE agreement 0.83 in four cell lines; Supplementary Fig. S11h, i), DDX55 and PABPN1 peaks lay on the structure; neither protein bound any of the five same-gene controls, and each had its strongest peak in the gene on this structure. The structure lies 64–165 nt upstream of the gene’s main polyA site in PolyASite 2.0,^57^ between the UGUA motif bound by CFIm and the AAUAAA hexamer. In a proposed model, such a fold between the two motifs brings them together to promote use of the site, and DDX55 represses use of the site by unwinding the fold;^58^ PABPN1 also suppresses alternative polyA sites.^59^ The *HMGCS1* structure may thus promote use of this site, restrained by DDX55. RLMFam11457 has a high scan lfdr (0.86) but is aligned across 38 species, agrees with in-cell reactivity and was also predicted by the RNAz and CMfinder screens.^13,15^

Supplementary Fig. S11a–e shows the pri-miR-22 positive control (a sharp DGCR8 signal on its predicted 15-base-pair helix), the *PTPRU* 3′UTR (RLMFam03987; as in *NUTF2*, a Staufen2 peak on the 5′ arm of an 11-base-pair helix, with UPF1 bound across the 3′UTR) and three candidates in which a protein that recognizes RNA structure binds the predicted helix. In the *SLC39A9* 3′UTR (RLMFam02146), LIN28B, which binds many mRNAs and regulates their translation,^60,61^ bound a 16-base-pair helix in both cell lines. In an intron of *MACF1* (RLMFam04531), DGCR8, which also binds hairpins in mRNAs outside annotated miRNA genes and directs their cleavage by the Microprocessor,^62,63^ bound a hairpin with a 20-base-pair stem, with no annotated miRNA nearby. In the first intron of the main *CADM1* transcript, part of the 5′UTR of a minor transcript (RLM-Fam00511), exportin-5 (XPO5), which also binds and regulates the *DICER1* mRNA,^64^ bound a long imperfect hairpin. LIN28B and DGCR8 were also among the RBPs found more often on candidates than on low-score regions, at unadjusted P < 0.05 (exploratory; no RBP passed BH q < 0.05; Fig. 6b). These pairings suggest that LIN28B may regulate the *SLC39A9* (ZIP9) mRNA through its 3′UTR, that the Microprocessor may cleave the *MACF1* pre-mRNA at the intronic hairpin and that exportin-5 may bind *CADM1* transcripts through this hairpin. Supplementary Table S6 lists 21 such hand-selected pairings (seven are known elements; the list is illustrative, not chosen by fixed criteria) with their evidence and literature support. Together, these cases show how RLM-Fam leads to hypotheses about the functions of individual RNAs, each to be tested with structure-disrupting and compensatory mutations.

### Structures with higher RLMHelix scores recruit structure-recognizing proteins in pull-down experiments

The ENCODE eCLIP data cover a fixed panel of 168 RBPs in two cell lines. To test, without preselecting proteins, whether the scores relate to the recovery of structure-recognizing binders, we used pull-down mass spectrometry (Fig. 6d–f) with 24 RNAs chosen independently of this scan from conserved structures predicted by comparative genomics (EvoFam^14^ and Hou et al.^65^), including let-7, the *MAT2A* hairpins and SNORA2 (Supplementary Table S7); the binders of each bait were identified by comparison with the other 23 (Supplementary Fig. S12). Twelve of the 16 human baits of at least 30 nt with alignments recruited a structure-recognizing binder, either a dsRBP, which reads duplexes, or a known partner that reads a sequence in a hairpin loop (LIN28B^66^ for let-7 and METTL16^42^ for the *MAT2A* hairpins; Fig. 6e).

Baits that recruited a structure-recognizing binder scored higher than the four that did not. With each score expressed as the distance from its shuffles (z), the AUC was 0.92 for RLMHelix-MSA (P = 0.013; Fig. 6f); single-sequence RLMHelix showed the same trend but, with 16 baits, did not reach significance (AUC 0.81, P = 0.078; Supplementary Fig. S13c), and thermodynamic scores (RNAalifold −MFE, RNAfold −MFE and RNAz) reached only 0.52–0.56. In paired comparisons on the same baits, RLMHelix exceeded −MFE both with alignments (RNAalifold) and on single sequences (RNAfold), whereas the difference from RNAz was not significant (P = 0.061; Fig. 6f, Supplementary Fig. S13c). The z of RLMHelix-MSA also correlated with the enrichment of all quantified dsRBPs (ρ = 0.52, P = 0.037), more strongly than with that of all RBPs (ρ = 0.31; the intervals overlap; Supplementary Fig. S13b, d). Although the baits were few, recovery of a structure-recognizing binder was associated with the score of helices supported by dependencies rather than with thermodynamic stability. Moreover, the baits came from earlier comparative screens rather than from RLM-Fam. Helices supported by language-model dependencies may therefore capture structural features that RBPs recognize, which would make the score a guide to functional RNA structures beyond those catalogued here.

## Discussion

This study implemented, for the genomes of humans and other vertebrates, a search that reverses the order of comparative genomics: RLMHelix proposes structural hypotheses from single sequences, and alignment starts from the regions that carry them. Distillation reduced a scan of the transcribed human genome from hundreds of GPU-days to a few days on a single desktop GPU (NVIDIA GB10). Two-thirds of the selected candidates were new to four earlier genome-wide screens, yet their predicted structures agreed with in-cell reactivity, and more than half of the candidates tested kept their structural signal in homologues aligned without assuming a structure. Higher scores accompanied more binding by double-stranded RNA-binding proteins and, in exploratory pull-downs, recovery of structure-recognizing proteins.

Discovering new RNA structures by scanning genomes with language models applied to single sequences has so far been demonstrated in prokaryotes, from dependencies^27^ or from attention maps.^67^ The genomes of humans and other vertebrates are two to three orders of magnitude larger, structured RNAs are sparse and base composition varies widely; ranked by the raw score, AT-rich windows came to the top of our scan. Discovery in these genomes therefore requires composition-aware calibration, estimates of false discovery and filtering by conservation and structural evidence, which we combined into a single procedure. Once calibrated, dependencies added information that thermodynamics lacks. In the pooled Rfam benchmark, the known RNAs at the top of the composition-calibrated ScanFold 2.0 ranking were almost all miRNA precursors, which are unusually stable for their base composition, whereas RLMHelix also ranked snoRNAs highly; in the genome scan, RLMHelix enriched known RNAs among the top windows about twice as strongly as ScanFold 2.0.

This study has three main limitations. First, the score assumes pseudoknot-free structures and continuous stems, so it is poorly suited to pseudoknots and structures broken by bulges, and it did not distinguish long RNAs from their shuffles (Fig. 2f). Because the scan used genomic sequence, structures that span exon junctions were also outside its reach. Absence from RLM-Fam therefore does not mean absence of structure. Second, the calibration was built for 200-nt genomic windows, so short sequences would need calibrations of their own, as would large sets of alignments ranked against one another with RLMHelix-MSA. The catalogue was not selected at a fixed false discovery rate. Against SISSIz shuffles, the background on which earlier screens based their false discovery rates, an estimated 13% of candidates had homologue alignments indistinguishable from their shuffles, close to the 14% reported for the CMfinder screen.^15^ Alignments of low-score regions in the same genes, however, also exceeded their shuffles more often than expected; with these as the background, the estimate rises to 25%, and the shuffle-based estimates of earlier screens are probably low for the same reason. Each candidate therefore carries its lfdr and evidence tier (Supplementary Table S4), so that users can choose the support their experiments require.

Third, the evidence on protein binding is limited by the available data. Public eCLIP data cover 168 RBPs in two cell lines, and about 70% of partners assigned from eCLIP alone could arise by chance. Combining binding with the properties of the partner, conservation and known examples nonetheless yielded testable hypotheses, such as LIN28B on a helix in the *SLC39A9* 3′UTR and DGCR8 on an intronic hairpin of *MACF1* (Fig. 6b; Supplementary Fig. S11c, d); binding maps for more proteins and cell types would extend such assignments to many more candidates. The pull-down experiment was a small, exploratory analysis of existing conserved structures, and baits drawn from RLM-Fam would test the scan directly.

RLM-Fam is intended to supply candidates for experiments on unknown RNA functions. Grouping into a family does not guarantee shared function,^16^ and as the *MAT2A* hairpins were found by a comparative screen and later linked to regulation through METTL16, linking structure to function requires experiments.^14,42^ Structure-disrupting and compensatory mutations in structures such as those in *NUTF2, HMGCS1, SLC39A9, MACF1* and *CADM1* would test their proposed effects on binding, mRNA decay, polyA-site use, translation and pre-mRNA processing. Human candidates with structural and functional support are targets for discovering new regulatory mechanisms and, like the miR-96 precursor and the *MALAT1* triple helix, may also be targets for small molecules that act on RNA structure.^7,8^ More broadly, reversing the order of comparative genomics brings parts of the human genome where genome alignments are unreliable within reach of the search for conserved RNA structures. As RNA language models improve and binding and probing data cover more proteins and cell types, the same approach could extend to other genomes and transcript isoforms and help to map the functional RNA structures that matter for human health and disease.

## Methods

Details of each section, the measurement of runtime, the training of the student and the full pull-down protocol are given in the Supplementary Methods.

### Dependency maps

Dependency maps (the teacher) were computed with the RNA language model RiNALMo-giga (Hugging Face model multimolecule/rinalmo-giga).^28^ The reference sequence was input without masking, and nucleotide predictions at all positions were obtained for the reference and for every variant in which a single position was replaced by one of the three other nucleotides. For a sequence *x* of length *L*, let *p*_*j*_(*k* ∣ *x*) be the probability that the model assigns to nucleotide *k* at position *j*, and let *x*^*i*→*a*^ be *x* with position *i* replaced by nucleotide *a* ≠ *x*_*i*_. The dependency of position *j* on position *i* is

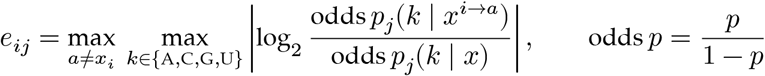

with *e*_*ii*_ = 0 and probabilities clipped to [10^−7^, 1 − 10^−7^].^27^ In words, *e*_*ij*_ is the largest change, in log2 odds, of any nucleotide prediction at *j* when the base at *i* is substituted. A sequence of length *L* therefore requires 3*L* + 1 forward passes. Directional dependencies were symmetrized as *D*_*ij*_ = max(*e*_*ij*_, *e*_*ji*_) and corrected for background (next section).

### Background correction by APC

The average product correction (APC) estimates a background component from the mean interaction strength of each position and subtracts it; it was proposed for mutual information in protein sequences.^32^ We applied the same correction to the symmetrized dependency map *D*. With *V*_*i*_ = {*j* : |*i* − *j*| ≥ 4}, 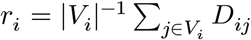 and *g* the mean of *D*_*ij*_ over all position pairs with |*i* − *j*| ≥ 4, the teacher target was

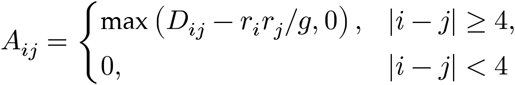

When *g* = 0, no background is subtracted. APC here is a background correction of dependencies, not a statistical test for evolutionary covariation between aligned sequences.

### Student distillation

Computing a dependency map takes 3*L*+1 forward passes of the language model, too many for a genome scan. We therefore used knowledge distillation, in which a small, fast model (the student) is trained to reproduce the outputs of a slower one (the teacher; here, the APC-corrected dependency maps).^34^ The student is a bilinear head (2.2 million trainable parameters) on the final-layer representations of the same frozen backbone, so that a single forward pass gives the APC-corrected dependency map Â. The 1,280-dimensional representation of each position is layer-normalized and mapped by two separately parameterized towers (Linear(1280, 768), SiLU, dropout 0.1, Linear(768, 128), SiLU) to source and target vectors *S*_*i*_, *T*_*i*_ ∈ ℝ^128^. With 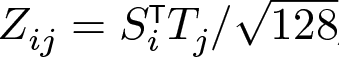, the predicted map is Â _*ij*_ = softplus[(*Z*_*ij*_ + *Z* _*ji*_)/2], with the diagonal set to 0. The student was trained to reproduce the teacher’s APC maps on synthetic sequences of 30–300 nt, drawn uniformly or with composition varied by a Dirichlet distribution, with no genomic or Rfam sequences, using a SmoothL1 loss that up-weights the top 5% and 1% of teacher values. The epoch was chosen by the pixel-wise Pearson correlation of APC maps on the validation set (0.512).

### Base-pair weights, dynamic programming and the RLMHelix score

Single-sequence base-pair probabilities (BPP) and minimum free energy (MFE) structures were computed with RNAfold from ViennaRNA (37 °C, default parameters).^68,69^ With the predicted APC dependency Â _*ij*_ and the BPP *P*_*ij*_, the weight of a base pair is defined as

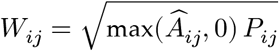

The weight is small whenever either the dependency or the BPP is low, so a pair contributes much only when both support it. A Nussinov-type dynamic programming (DP) algorithm chose the set of non-crossing base pairs, with each nucleotide in at most one pair, that maximizes the sum of weights.^33^ Allowed pairs were AU, GC and GU, and the minimum loop length was 3. The quantity optimized is the total weight, not the number of pairs or the free energy. After decoding, isolated pairs without an adjacent stacked pair were removed. A stem is a run of consecutive pairs (*i, j*), (*i* + 1, *j* − 1), …, and the RLMHelix score is the total weight of stems of at least four consecutive pairs (minL4); the minimum stem length is applied after decoding, not inside the DP, and the score is not normalized by length or number of stems. Combining pairwise evidence with BPP also underlies CMfinder and RNAPhylo; here, however, the geometric mean of the two is used as a DP weight rather than to modify emission probabilities within a model.^12,39^

### MSA score (RLMHelix-MSA)

RLMHelix-MSA maps the APC dependencies predicted by the student for each sequence of an MSA onto alignment coordinates, averages them per position pair and applies the same DP and stem scoring as for single sequences. No additional training for discriminating structured RNAs was performed. In each window of 200 columns of the reference (RF) annotation of the alignment, moved in steps of 100 columns, the alignment was pruned to six sequences, always keeping the human row, with the algorithm of RNAz’s rnazSelectSeqs.pl (one sequence of any pair with more than 0.99 identity is removed, and the sequence with the largest sum of squared deviations from 0.80 identity is then removed repeatedly), and sequences were limited to ungapped lengths of 30–300 nt. Position pairs covered by fewer than half of the sequences were set to 0. The weight is the geometric mean of the averaged dependency and the RNAalifold BPP; using the averaged dependency alone gives the ‘dependency only’ variant. The MSA score is the maximum over windows.

### Benchmark sets, baselines and metrics

#### Rfam

From Rfam 15.1 vertebrate alignments mapped to the assemblies of the UCSC hg38 100-way alignment, one human representative sequence was chosen per family.^41^ The 328 families whose representative consists of ACGU only and is 30–300 nt long, whose alignment includes at least four assemblies and that are shared with the MSA set were fixed as the common benchmark for single-sequence, MSA and phylogenetic-model comparisons (Supplementary Table S1). Single-sequence negatives were 100 dinucleotide-preserving shuffles of each representative (uShuffle, k = 2; 33,128 sequences in total with the representatives).^36^ MSA evaluation used 322 families, leaving out six miRNA families whose alignments collapse to a single sequence on pruning, with 100 negatives per family simulated by SISSIz (`--simulate -d`), which preserves phylogeny and composition.^37^

#### EvoFam and RNAz 2005

MSAs were cut from the UCSC hg38 100-way alignment projected onto human coordinates for EvoFam loci (the genome-wide (GW) set of Parker et al., 725 EvoFold v2 predictions in 220 families; 318 loci in 122 families were scored by every method)^14^ and for 598 regions sampled from the predictions of the RNAz screen of the human genome,^13^ each with 100 SISSIz negatives. The RNAz 2005 regions are not a ground-truth set; RNAz itself selected them, so comparisons with RNAz are biased in its favour. Computations that failed were recorded with their causes and never scored as 0.

#### Baselines

Single-sequence baselines were the RNAfold MFE and MFE z-scores calibrated for sequence composition with ScanFold 2.0 (mononucleotide- and dinucleotide-shuffle models)^25^ and with RNAz 2.1 regression.^38^ MSA baselines were the RNAalifold free energy (−MFE),^70^ a BPP-only score, RNAz (the maximum RNA-class probability over windows)^10^ and two phylogenetic models, RNAPhylo^39^ and EvoFold.^11^ The ablations ‘dependency only’ and ‘BPP only’ used the same DP and stem sum as RLMHelix, with the pair weight replaced by the dependency or by the BPP. RNAPhylo inferred structures with the Cocke–Younger–Kasami (CYK) algorithm and was scored with the phylo-SCFG posteriors alone and, as in the published pscore of CMfinder, with the posteriors weighted by the thermodynamic partition function.^15^ Both settings are called RNAPhylo below, the second without qualification. The phylogenetic models scored whole alignments with the UCSC hg38 100-way tree, whereas the other MSA scores were computed on the same 200-column windows of pruned alignments (Supplementary Methods).

#### Metrics

Each known RNA (or MSA) was compared with its own 100 shuffles to obtain the area under the precision–recall curve (AUPRC; average precision) and the empirical P value (1+#{*s*_null_ ≥ *s*_native_})/101. With a single positive, the expected AUPRC of a random ranking is *H*_101_/101 = 0.051, where 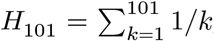; the mean AUPRC is the unweighted mean across families (macro_auprc in Supplementary Table S2; for EvoFam, loci were first averaged within families). To mimic scanning, all sequences were also ranked together to obtain pooled precision–recall curves and the average precision (AP), which is approximately the fraction of positives (1/101) for a random ranking. Intervals and two-sided P values for paired differences are from paired bootstraps over families (over regions for RNAz 2005), without correction for multiple comparisons. Because shuffles do not carry the base pairs of the original sequence, base-pair-level AUROCs were not computed for negatives.

### Genome scan, calibration and false discovery

#### Scan

Positions of hg38 in the introns and UTRs of GENCODE v50 mRNAs and within lncRNA genes were scanned on the strand of the annotation in 200-nt windows with a 100-nt step.^71^ Windows containing repeats (RepeatMasker, tantan for tandem repeats, Inverted Repeats Finder for inverted repeats; thresholds in the Supplementary Methods) or lying in intergenic regions were removed,^72,73^ leaving 7,869,821 windows (6,924,574 distinct coordinates), each scored from the dependencies of the student and the RNAfold BPP of the window.

#### Calibration

To rank windows of different lengths and compositions together, we computed a P value for each window with a conditional generalized extreme value (GEV) model of the null distribution of log(1+*s*), whose location and scale depend on 25 covariates of length and mono- and dinucleotide composition and whose shape is shared. It was fitted, separately from student training, to 570,000 scores of dinucleotide-preserving shuffles of uniform, Dirichlet and genomic sequences. The Rfam benchmark used raw scores, because each known RNA is compared with its own shuffles and the calibration, built for 200-nt windows, is anti-conservative for short sequences: 11.6% of the Rfam shuffles reached a nominal P ≤ 0.05, mainly at 30–99 nt. The Benjamini–Hochberg (BH) procedure across all 7,869,821 windows defined, at a nominal q ≤ 0.05, a stricter reference set;^74^ its nominal level was not verified for overlapping windows, merging and later selection.

#### Local false discovery rate

Because calibrated P values were conservative overall on real windows, we estimated an empirical null.^40^ P values were transformed to *z* = −Φ^−1^(*p*); the null *N*(*μ*_0_, 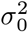) and the null proportion *π*_0_ were estimated by central matching (*μ*_0_ = 0.673, *σ*_0_ = 0.548, *π*_0_ = 0.951), and the local false discovery rate lfdr(*z*) = *π*_0_*f*_0_(*z*)/*f*(*z*) was computed with the mixture density *f* estimated by Lindsey’s method. Estimates beyond the top 0.1% depend on extrapolating the null into the tail, and the lfdr does not reflect the later lineage conditions.

### Candidate selection and alignments

#### Stage 1

Structural cores were extracted from the top 1% of windows by calibrated P value (78,699 windows; P ≤ 0.02088). After stems of fewer than four pairs were removed from the DP decoding of each window, each outermost base-paired interval became a core, as in EvoFold.^11^ Overlapping top windows on the same strand were merged into loci, each with one primary core and a peak P value (the smallest calibrated P value among its windows), and identical cores on either strand were collapsed (71,196 representatives).

#### Stage 2

Each core was searched with Riboseek against the genomes of 20 vertebrates (9 non-human primates, 4 non-primate mammals, 3 non-mammalian tetrapods and 4 other vertebrates; Ensembl release 116 assemblies, listed in Supplementary Table S8).^75^ Hits with E ≤ 10^−3^ and query coverage ≥ 0.8 were high-confidence hits, and cores meeting any of four routes were selected (15,913):

- vertebrate_wide: ≥ 1 non-mammalian tetrapod and ≥ 1 other vertebrate
- mammal_conserved: ≥ 3 non-primate mammals and ≥ 3 primates
- primate_focused: ≥ 6 primates and peak P ≤ 0.001
- high_score_rescue: peak P ≤ 0.0005 (no lineage condition)

The route names are selection strata of the search; they do not imply a lineage-specific origin.

#### Alignments

Seed alignments were built from high-confidence hits, and covariance models (CMs) were built from them with the human structure as the consensus structure (SS_cons; Infernal).^76^ The alignments were extended with homologues from 34 further vertebrates and from paralogous loci in the human genome and realigned with the CM, yielding alignments of up to 55 species (final55; median 34) for 15,821 candidates (the RLM-Fam candidates). Provisional families (14,037) were connected components of candidates whose seed CMs scored each other’s homologues above a relative threshold.

### Comparison with earlier screens and with thermodynamic scanning

#### Earlier screens

Candidates were compared with the predictions of four earlier genome-wide screens lifted to hg38: RNAz 2005,^13^ EvoFold (original and v2, the predictions from which the EvoFam families were built)^11,14^ and the CMfinder CRSs of Seemann 2017 (all published and those with a reported FDR ≤ 10%).^15^ Overlaps of at least 1 bp on either strand were counted, and the background was the overlap of intervals of the same lengths placed at random within the scanned space (`bedtools shuffle`).^77^ The input regions of the RNAz screen are no longer distributed and were approximated by PhastCons conserved elements, which EvoFold also scanned.^11,48^

#### Rfam enrichment along the selection

At each stage, enrichment was the fraction of intervals overlapping significant Rfam 15.1 hits on the same strand, divided by that fraction for intervals of the same lengths placed at random as above.^41^

#### Thermodynamic scanning on the scan windows (Fig. 4e, f)

All 7,869,821 windows were ranked by calibrated RLMHelix P value, RNAfold MFE and ScanFold 2.0 z-scores, with the 4,575 windows overlapping a significant Rfam hit on the same strand as positives; intervals are from a block bootstrap over 1-Mb blocks.

### Structural support

#### In-cell reactivity

We used published processed icSHAPE reactivities of six human cell lines (GSE145805 and GSE74353).^49,50,78^ For cores measured over at least 50% of their length, the AUROC of reactivity for unpaired versus paired nucleotides was compared with that of the same labels circularly shifted by every possible offset. Cores were the unit of inference for the comparisons with the shifted control and between structure methods, and RLM-Fam candidates for the comparison between groups (Fig. 5a, right); candidates with an empirical P ≤ 0.05 in at least half of their cell lines were counted as supported by icSHAPE.

#### Covariation

Covariation of the proposed base pairs was tested with R-scape 2.6.16 under three alignment conditions:^20,51^ (i) the final55 CM alignment (CM-aligned), (ii) the same sequences realigned with MAFFT (sequence-realigned) and (iii) Riboseek high-confidence hits aligned with MAFFT without structural information (structure-free).^52^ We took condition (iii), which depends neither on CM-based selection of homologues nor on CM alignment, as primary. The rate of chance positives was estimated by applying R-scape with the same structure to one SISSIz shuffle of each alignment of conditions (ii) and (iii).^37^ We never rejected a structure solely because its covariation was not significant. Consensus structures with significantly covarying pairs were drawn from the R-scape output with R2R (the version distributed with R-scape 2.6.16).^53^

#### Structural signal in homologues

Each alignment of condition (iii) was scored with RLMHelix-MSA under the settings of Fig. 3 together with 19 SISSIz shuffles. Because the human sequence was chosen by the scan score, the human row was removed before scoring. An alignment scoring above all 19 shuffles (empirical P = 1/20) kept the signal, so the chance rate is 5%; the 14,565 candidates for which all 19 shuffles were scored were analysed. The false discovery rate of the catalogue was estimated with Storey’s method from the ranks of the alignments among their shuffles, which are uniform for an alignment without structure: *π*_0_ = *P* (rank ≥ 11)/0.5.^79^ To check this background on real alignments, 600 randomly chosen candidates and, for each, a low-score region in the same gene (matched for length, region type and GC; the controls of the eCLIP analysis) were searched, aligned and scored in the same way.

### Comparison with RNA-binding protein binding (eCLIP)

#### Data and matched controls

Reproducible eCLIP peaks from ENCODE (K562 and HepG2; 250 experiments, 168 RBPs) were overlapped on the same strand with the candidate cores, control intervals and scan windows described below.^54,80^ To separate binding to candidates from binding to where candidates lie, each candidate core was matched with up to five control intervals in the same gene, with the same length, region type and exon status, the closest GC content and a low score (below the genome-wide median and calibrated P > 0.1). For each experiment, a Mantel–Haenszel odds ratio and a Cochran–Mantel–Haenszel test were computed with each candidate and its controls as one stratum, and P values were BH-adjusted across experiments (Supplementary Table S5).^81^ For comparison, an analysis following Seemann 2017 used all scan windows as background.^15^

#### Dose–response and double-stranded RNA-binding proteins

For the dose–response, window-level odds ratios were computed in bins of calibrated P value, stratified by gene, region type, exon overlap and GC. We defined six RBPs that recognize double-stranded RNA (dsRBPs; DGCR8, DROSHA, ILF3, STAU2, DHX30 and XPO5) as a group before examining per-RBP results, and also performed an analysis excluding candidates within 500 nt of a miRNA gene.

#### Candidate-specific RBPs and the chance ratio

An RBP was called candidate-specific when its peak overlapped the core, none of the candidate’s controls was bound and the peak was the strongest of that experiment in the host gene. The chance ratio, the number of pseudo-candidates with a candidate-specific RBP divided by the number of candidates with one, was estimated by treating one control as a pseudo-candidate and repeating the calls with the remaining controls; for each RBP, calls at candidates and at pseudo-candidates were compared with McNemar’s test.^82^

### Known functional structures

Known functional cis-acting structures in human mRNAs and lncRNAs (82 elements of 12 classes from 31 papers; Supplementary Table S3) were compared with length-matched intervals in the same genes.

### Evidence tiers

Each catalogue candidate was annotated with statistical support (lfdr ≤ 0.2, which all candidates containing a BH-accepted window meet) and independent structural support (icSHAPE support or covariation under condition (iii)). Candidates with both were assigned to tier A, those with one to tier B and those with neither to tier C (Supplementary Table S4). We did not treat absent covariation or missing icSHAPE measurements as evidence against a candidate. We also selected by hand 21 pairings of a candidate structure with a partner, with grounds from eCLIP or from literature-defined structure–sequence rules (for the targets of METTL16, SLBP and iron regulatory proteins, and for long double-stranded RNA), and listed them with their literature support (Supplementary Table S6).

### Pull-down mass spectrometry (Fig. 6d–f, Supplementary Figs. S12 and S13)

#### Baits and pull-down

The 24 baits, chosen independently of this scan, were 16 EvoFam elements, including let-7, the *MAT2A* hairpins and SNORA2,^14^ and 8 conserved structures found with CMfinder by Hou et al. (hereafter CM-line; 6 human, 2 mouse)^65^ (Supplementary Table S7). Bait RNAs were transcribed in vitro with the MEGAshortscript T7 Transcription Kit (Thermo Fisher Scientific), conjugated with FLAG at the 3′ end by periodate oxidation and FLAG-hydrazide coupling, and incubated with 293T cell lysate, anti-FLAG M2 antibody and Protein G magnetic beads. Bound complexes were eluted with FLAG peptide, processed by the SP3 method^83^ and analysed on an Evosep One system coupled to an Orbitrap Astral mass spectrometer in data-independent acquisition mode. Each bait was pulled down twice, and each pull-down was measured twice (four runs per bait, 96 runs in total). Proteins were identified and quantified with DIA-NN 2.0.2 (1% false discovery rate) and its MaxLFQ algorithm.^84,85^

#### Binder calls and labels

MaxLFQ intensities were log2-transformed and median-normalized per sample, a proDA model with a mean per bait and an effect of the pull-down replicate was fitted to the 96 runs,^86^ and each bait was compared with the mean of the other 23 by a Wald test. We took proteins with adjusted P < 0.05 and log2 fold change ≥ 2 as binders of that bait. A bait had a structure-recognizing binder when its binders included a double-stranded RNA-binding protein (17 proteins listed in the Supplementary Methods or proteins with a Pfam double-stranded RNA-binding domain), which reads duplexes, or a known partner that reads a sequence in a hairpin loop (LIN28A and LIN28B for let-7; METTL16 for the *MAT2A* hairpins; DKC1, NOP10, NHP2 and GAR1 for SNORA2; DGCR8 and DROSHA for miR-9-2; and SLBP for the histone stem-loop).

#### Comparison with the scores

The 16 human baits of at least 30 nt for which alignments could be built (10 EvoFam, 6 CM-line) were scored with the methods of Figs. 2 and 3 against 100 shuffles (SISSIz for MSAs, uShuffle for single sequences),^36,37^ and each score was converted to z = (score − mean of shuffles) / s.d. of the shuffles of that bait. We computed ROC curves and the AUC for baits with versus without a structure-recognizing binder (two-sided Mann–Whitney tests; paired DeLong tests between scores)^87^ and the Spearman correlation of z with the number of double-stranded RNA-binding proteins among the binders. Without a threshold, z was also correlated with the mean log2 fold change, against the other 23 baits, of all quantified double-stranded RNA-binding proteins (and, as a control, of all quantified RNA-binding proteins). All 95% intervals are from 10,000 bootstrap resamples of baits.

### Statistics

The unit of inference was the family for the Rfam and EvoFam benchmarks and the region for RNAz 2005, the 1-Mb block for comparisons over scan windows, the core for the icSHAPE comparisons with controls and between structure methods, the candidate for icSHAPE support, covariation and the structural signal in homologues, the experiment for eCLIP enrichment and the bait for the pull-down. Intervals are from paired bootstraps over families (or regions) for paired differences on the benchmarks, from a block bootstrap over 1-Mb blocks for scan windows and from 10,000 bootstrap resamples of baits for the pull-down. P values for paired differences on the benchmarks are two-sided bootstrap values without correction for multiple comparisons. eCLIP enrichment was tested per experiment with Cochran–Mantel–Haenszel tests and BH-adjusted across experiments. Candidate-specific RBP calls were compared with McNemar’s tests, and the pull-down used two-sided Mann–Whitney tests and paired DeLong tests between scores. Empirical P values from shuffles or circular shifts are described in the sections that use them. Random seeds and the tests of analyses not listed here are given in the Supplementary Methods.

### Use of large language models

Large language models (Claude, Anthropic; Codex, OpenAI) were used to assist in writing the analysis code and the manuscript text. All outputs were reviewed, tested and edited by the authors, who take full responsibility for the content.

Data and code availability are given in the sections below.

#### Reporting summary

Further information on research design is available in the Nature Portfolio Reporting Summary linked to this article.

## Data availability

The RLM-Fam catalogue (candidate table, alignments, covariance models and provisional families) and the coordinates, RLMHelix scores, calibrated P values and BH q values of all 7.87 million scan windows (about 280 MB) will be deposited on Zenodo and made public upon acceptance; the candidate and family tables are provided as Supplementary Table S4. The raw data and the protein identification and quantification results of the pull-down mass spectrometry will be deposited to the ProteomeXchange Consortium via the PRIDE repository and made public upon acceptance. All other data analysed in this study are public; sources and versions are given in the Methods and Supplementary Table S8.

## Code availability

The RLMHelix code, the trained student head and the score calibrator are available under the MIT License at https://github.com/Shunsuke-1994/rlmhelix (version 0.1.0), with documentation, example data and tests; the version used in this study will be archived on Zenodo upon publication.

## Acknowledgements

This work was supported by JST ACT-X (JPMJAX24L8 to S.S.), JSPS KAKENHI (JP24K23175 and JP26K18281 to S.S.) and JST CREST (JPMJCR23B3 to H.S., M.H. and S.A.).

## Author contributions

S.S. conceived the study, developed RLMHelix and performed the computational analyses. S.A., K.M. and M.O. performed the pull-down mass spectrometry. H.S. and M.H. supervised the project. S.S., H.S. and M.H. wrote the manuscript. All authors reviewed the manuscript.

## Additional information

Supplementary Information is available for this paper.

