## Supplementary Information for "Alignment-free discovery of conserved RNA structures in the human genome"

### Supplementary Methods

These methods give the details behind the sections of the Methods. The definitions of the dependency map, APC, the RLMHelix score and RLMHelix-MSA are given in full in the Methods and are not repeated.

#### Student architecture and training

The student is a bilinear head that outputs the APC-corrected dependency map directly from a single forward pass of the same frozen backbone. The 1,280-dimensional representation of each position was layer-normalized and passed to separately parameterized source and target towers (Linear(1280, 768), SiLU, dropout 0.1, Linear(768, 128), SiLU). With outputs $S_i, T_i \in \mathbb{R}^{128}$, $Z_{ij} = S_i^{\top} T_j / \sqrt{128}$ and $\widehat{A}_{ij} = \mathrm{softplus}[(Z_{ij} + Z_{ji})/2]$, with the diagonal set to 0. The head has 2,167,040 trainable parameters, excluding the frozen backbone. The teacher is symmetrized by the directional maximum and the student by the directional mean. APC is not reapplied to the student output.

Training sequences came from two synthetic generators and included no genomic or Rfam sequences. One draws each nucleotide uniformly (uniform; 27,000 training and 3,000 validation sequences); the other varies the composition with a Dirichlet distribution (alpha 3, 5, 10, 20, 50 and 100; 32,400 training and 3,600 validation sequences). Lengths were 30–300 nt, and each generator was sampled with probability 0.5 during training. With AdamW (learning rate $2 \times 10^{-4}$, weight decay $10^{-4}$), we minimized a tail-weighted SmoothL1 loss against the teacher APC values (negative values set to 0). The loss covered upper-triangular position pairs with $|i-j| \geq 4$: pairs in the top 5% of teacher values received weight 3, those in the top 1% weight 10 (replacing, not adding to, 3) and all others weight 1. It was normalized by the sum of weights within each sequence and then averaged across sequences. No log transform, z-scoring, upper clipping or length normalization was applied. We selected epoch 13, which maximized the pixel-wise Pearson correlation of APC maps on the validation set (0.512). This correlation is a different measure from base-pair F1 or the detection rate of conserved structures.

#### Runtime measurement

Teacher and student runtimes were measured on the same GPU (NVIDIA GB10, bf16 autocast, SDPA attention) (Fig. 1d, e). Inputs were 20 sequences per length drawn from chr22 scan windows (transcribed non-coding regions, 200 nt), at lengths of 50, 100, 200 and 300 nt (300-nt inputs joined adjacent windows). The teacher ran the 3L + 1 sequences (the reference and its 3L single-nucleotide substitutions) in batches; the student was timed one sequence at a time and with 64 sequences per batch. DP and stem scoring are identical for the two and were timed separately on CPU. Genome-wide compute time was estimated by multiplying the 200-nt median by all 7,869,821 scan windows; for the student, the estimate is the sum of batched GPU inference and DP on 18 parallel CPU workers.

#### Benchmark sets

**Rfam.** Benchmark sets were built from Rfam 15.1 vertebrate alignments mapped to the assemblies of the UCSC hg38 100-way alignment.[41] For single-sequence evaluation, one representative sequence was chosen per family from its hg38 rows. After collapsing identical human sequences, the representative was chosen by the fewest non-ACGU characters, then coverage of consensus columns, sequence length and sequence ID. Of the 639 families with a representative consisting of ACGU only, 30–300 nt long and with at least four assemblies, the 328 shared with the MSA set (families with one sequence per assembly after identical sequences are collapsed) were fixed as the common benchmark for single-sequence, MSA and phylogenetic-model comparisons (Supplementary Table S1). Negatives were 100 dinucleotide-preserving shuffles of each representative generated with uShuffle ($k = 2$) (33,128 sequences including the 328 representatives; dinucleotide counts were checked against the originals).[36]

MSA evaluation used 322 families: the 328 minus six miRNA families (RF00687, RF00707, RF00741, RF00771, RF00870, RF00877) whose alignments collapse to a single sequence under RNAz-style pruning (Methods, RLMHelix-MSA) and therefore carry no information for any MSA method. Negatives were 100 alignments simulated with SISSIz (`--simulate -d -n 100`), which preserves phylogeny and composition.[37] Negatives for which RNAz produced no window were dropped individually (57 negatives in 8 families).

**EvoFam.** Of the 725 sequences of the EvoFam GW set (220 families of EvoFold v2 predictions) lifted to hg38, the 722 on primary chromosomes were used, and MSAs were cut from the multiz100way alignment projected onto human coordinates.[14] Two predictions on hg18 `*_random` contigs that lift to the same hg38 interval as a primary-chromosome prediction of the same family were removed (720 loci). Non-ACGU characters were converted to gaps, and the human row and rows covering at least 50% of human columns were kept. For loci with a human length of 30–300 nt and at least four rows, 100 negatives were generated with SISSIz using the same settings as for Rfam. Evaluation used the 318 loci in 122 families for which the native MSA and all 100 shuffles were scored by every method (complete case).

**RNAz 2005 predictions.** From the predictions of the RNAz screen of the human genome (Set1, $P > 0.5$), we used the 6,498 regions that could be lifted to hg38.[13] We sampled 250 regions from each of four strata defined by RNAz confidence ($P > 0.9$ or not) and membership in Set3 (conserved to fish), in proportion to length bins (30–99, 100–149, 150–199 and 200–300 nt) within strata (seed 20260928). MSAs were built with the EvoFam rules; the first 600 regions in sampling order with an MSA and SISSIz shuffles were scored, and the 598 for which every method scored the native and all 100 shuffles were used. Because the predictions are unstranded, both strands were scored and each method took the larger score. These predictions are not a ground-truth set; RNAz itself selected them, so comparisons with RNAz are biased in its favour.

**Why the sets differ between methods.** The difference between single-sequence (328) and MSA (322) evaluation reflects the families whose alignments lose all information on pruning. For EvoFam and RNAz 2005, loci were limited to those for which MSAs could be built and every method returned a score. Failures were recorded with their causes and never scored as 0.

#### Baseline methods

Single-sequence baselines were the RNAfold minimum free energy (MFE) and MFE z-scores calibrated for sequence composition. ScanFold 2.0 z-scores were obtained with the published TensorFlow models, which predict the mean and standard deviation of shuffled-sequence MFEs from the same 20 sequence features as the ScanFold 2.0 code.[25] We used both the mononucleotide-shuffle (MeanMFE / StdDev) and dinucleotide-shuffle (DiMFE / DiStd) models at commit `5eec7c20a49b0a65b856069f34478e12415ad2b7`; these models were trained on sequences of 60–200 nt and were applied unchanged outside that range (30–59 and 201–300 nt). RNAz 2.1 regression z-scores were computed by applying `--no-shuffle -d` (dinucleotide background) to an alignment of two identical copies of each sequence, and sequences whose z-scores were computed outside the range covered by the regression were flagged.[38] Because negative z-scores indicate greater stability than expected, all z-scores were negated to serve as scores.

MSA baselines were the RNAalifold free energy (−MFE), a BPP-only score, RNAz and two phylogenetic models (RNAPhylo[39] and EvoFold[11]).[10,70] RNAz was run with `RNAz --forward` on windows of 200 RF columns moved in steps of 100, after keeping rows of 30–300 nt and pruning to six sequences (maximum identity 0.99, target identity 0.80). The MSA score was the maximum RNA-class probability over successful windows. RNAPhylo (the C implementation distributed with CMfinder 0.4.1.18) inferred structures by CYK without a given SS_cons and was scored in two settings: with the phylo-SCFG posteriors alone (no partition function) and, as in the published pscore, with the posteriors weighted by the thermodynamic partition function (`--partition`).[15] For the latter we built a version that applies the partition function only to the posterior, not to the CYK that infers the structure. RNAalifold (ViennaRNA Python API with default parameters), the BPP-only score and RLMHelix-MSA were computed, like RNAz, on the same windows of alignments pruned to six sequences, and the MSA score was the maximum over windows. The phylogenetic models instead scored whole alignments (all rows and columns, without windows or pruning), with gaps treated as missing data. Both used the UCSC hg38 100-way phylogeny (`hg38.100way.nh`) without pruning or rescaling; sequences were matched to its leaves by assembly name, and leaves without a sequence were marginalized. RNAPhylo was run as `RNAPhylo -t hg38.100way.nh -s single_model -p pair_model -g scfg --ignore-all-gap` (default gap threshold, 0.7), with `--partition` for the thermodynamic setting, and its score was the total pair posterior, the sum over CYK pairs of $-\log(1-\text{posterior})$. EvoFold was EvoFoldV2 of the phy library (commit `10ee240d`) with its distributed configuration (`EvoFoldV2 -c EvoFoldConfig.v7b`), and its score was the `score` field, the log ratio of the inside probabilities under the structural and the non-structural grammars, not `cykScore`. The phylogenetic scores of 271 of the 322 Rfam families were computed with an earlier macOS build of the same programs and those of the other 51 with a Linux build. For the phylogenetic models, a family was evaluated when its native MSA was scored and at least 50 valid negatives remained (invalid negatives were dropped individually, never set to 0).

#### Metrics

In per-family evaluation, each known RNA (or MSA) was compared with its own 100 shuffles to obtain the area under the precision–recall curve (AUPRC; average precision in scikit-learn) and the empirical P value $(1 + \#\{s_{\text{null}} \geq s_{\text{native}}\})/101$. With a single positive, the expected AUPRC of a random ranking is $H_{101}/101 = 0.051$. The unweighted mean across families is the macro AUPRC. For EvoFam, locus values were averaged within families, and families were then weighted equally. To mimic scanning, all sequences (33,128 for Rfam) were ranked together to obtain pooled precision–recall curves and the average precision (AP). The AP of a random ranking equals the fraction of positives (1/101). We also counted the known RNAs among the top k sequences (k = the number of known RNAs). Intervals are from paired bootstraps over families (over regions for RNAz 2005), without correction for multiple comparisons: for the MSA comparisons, 10,000 resamples for the macro AUPRC and 2,000 for differences in pooled AP (seed 20260909 for Rfam, 20260929 for the phylogenetic models on Rfam, and 20260928 for EvoFam and RNAz 2005), and, for the single-sequence comparisons, 2,000 for the macro AUPRC and 1,000 for differences in pooled AP. Two-sided P values for paired differences are twice the fraction of bootstrap differences on the smaller side of zero. Because shuffles do not carry the base pairs of the original sequence, base-pair-level AUROCs were not computed for negatives.

#### Genome scan

Positions of hg38 in the introns and UTRs of GENCODE v50 mRNAs and within lncRNA genes were scanned in 200-nt windows with a 100-nt step on the strand of the annotation (on both strands where both are annotated).[71] Because windows were placed with their centres inside these regions, some windows also overlap coding sequence. We removed windows with more than 40 nt (20% of the window) of RepeatMasker repeats (UCSC rmsk, all classes), with at least 20 nt of tandem repeats detected by tantan (`-f 3`), or containing an inverted repeat with an Inverted Repeats Finder (IRF) score of at least 40 (parameters `2 7 7 80 10 40 10000 200`).[72,73] Windows lying between genes were excluded, leaving 7,869,821 windows (6,924,574 distinct coordinates) for the analyses. For each window, the RLMHelix score was computed from the dependencies of the final student and the RNAfold BPP of the window.

**Score calibration and multiple testing**

To rank windows of different lengths and compositions together, we computed a P value for each window with a conditional generalized extreme value (GEV) model that predicts the null distribution of the RLMHelix score from sequence length and composition. Calibration was separate from student training and was fitted to $y = \log(1 + s)$. The fitting data were the scores of 570,000 dinucleotide-preserving shuffles (k = 2): 270,000 of uniform sequences, 270,000 of Dirichlet sequences (balanced to 45,000 per alpha) and 30,000 of 3,000 genomic prototypes. The 25 covariates were length L, log(1 + L), A/C/G/U frequencies, GC content, A/(A + U), C/(C + G) and the 16 dinucleotide frequencies, standardized with the means and standard deviations of the fitting set. The location was a linear function of the covariates, the scale the exponential of another linear function clipped to −5 to 8, and the shape a single $\xi = 0.4\tanh(\eta)$ shared by all sequences (estimate −0.1845). Upper-tail probabilities were computed with the SciPy GEV survival function. Short sequences with intermediate-to-high GC content and few CpGs are sparse in the fitting data, so calibrated P values tend to be too small for sequences like known Rfam RNAs.

The raw RLMHelix score and the calibrated P value have separate roles. The Rfam benchmark compares each known RNA with its own shuffles and therefore uses the raw RLMHelix score (the scan calibration is built for 200-nt genome windows and is anti-conservative for the short sequences of the Rfam set; see below); the genome scan uses calibrated P values. The Benjamini–Hochberg procedure was applied to the calibrated P values across all 7,869,821 eligible windows, and windows with a nominal q ≤ 0.05 formed a stricter reference set.[74] Windows overlapping on the same strand were merged into loci. We did not verify that this nominal level holds under the dependence between overlapping windows, the merging of windows into loci or the later selection steps. To check the calibration, we applied it to the 32,800 Rfam shuffles of Fig. 2: 11.6% reached a nominal P ≤ 0.05 and 0.8% P ≤ 0.001, with the excess mainly in sequences of 30–99 nt (14.1% and 1.1%; in the overlapping ranges of 100–199, 150–250 and 200–300 nt, 8.6% and 0.2%, 5.4% and 0.2%, and 3.4% and 0%).

**Empirical null and local false discovery rates**

Because calibrated P values were conservative overall on real windows, we estimated an empirical null in addition to the theoretical null used by the BH procedure.[40] P values were transformed to $z = -\Phi^{-1}(p)$, so that larger z indicates more structure than expected, and the null normal distribution $N(\mu_0, \sigma_0^2)$ was estimated by central matching: z values between the 25th and 75th percentiles were divided into 60 bins, and a quadratic was fitted to the log bin counts. $\pi_0$ was obtained from the null mass in that range ($\mu_0 = 0.673$, $\sigma_0 = 0.548$, $\pi_0 = 0.951$; Supplementary Fig. S4). The background fraction above a threshold z (Efron's tail-area false discovery rate) is $\mathrm{Fdr}(z) = \pi_0 S_0(z)/S(z)$, where $S_0$ and $S$ are the null and observed survival functions. The local false discovery rate (lfdr) is $\mathrm{lfdr}(z) = \pi_0 f_0(z)/f(z)$, with the mixture density $f$ estimated by Lindsey's method: histogram counts (bin width 0.05) were fitted by Poisson regression on a natural cubic spline with knots at quantiles of z. The lfdr was capped at 1 and made non-increasing in z to the right of $\mu_0$. $\mathrm{Fdr}(z)$ equals the mean lfdr above the threshold (Fig. 4c), and background fractions recomputed from the lfdr agreed with the direct estimate (0.612 and 0.613 for the top 1%; Supplementary Fig. S8). Both tails of z are heavier than normal, and estimates beyond the top 0.1% depend on extrapolating the normal fitted to the centre into the tail, so they are uncertain. The lfdr is derived from scan scores alone and does not reflect the later lineage conditions.

**Candidate selection and alignments**

**Stage 1.** Structural cores were extracted from the top 1% of windows by calibrated P value (78,699 windows; P ≤ 0.02088). In the DP decoding of each window, stems of fewer than four pairs were removed, and each outermost base-paired interval, over which the bracket depth rises from and returns to 0, became a core, as in EvoFold;[11] no flanking nucleotides were added. Top windows overlapping on the same strand were merged into loci, and in each locus the core with the largest total weight of retained pairs was the primary core. For each core, the pair of sequence and dot-bracket was compared with the pair of its reverse complement and reversed dot-bracket, and the lexicographically smaller pair was kept as the normalized orientation; identical normalized pairs were collapsed to one representative (71,196 representatives, chosen by ascending peak P value, the smallest calibrated P value among the windows of the locus, and then descending core score). For downstream analyses, representatives were restored to the sequence and structure of their source strand while keeping their IDs.

**Stage 2.** Each core was used as a query to search the genomes of 20 vertebrates, split into 10-kb chunks overlapping by 500 nt, with Riboseek (sensitivity 7.5, max-seqs 100, E ≤ 10).[75] The genomes of the panel (species and assemblies in Supplementary Table S8) were Ensembl release 116 assemblies, downloaded on 2026-08-26 as unmasked `dna` sequences: the primary assembly for human, mouse and zebrafish and the toplevel sequence otherwise, with sequences identical to release 115. For the selection routes below, the cores were searched in their normalized orientation (Stage 1); the seed alignments used a second search of the same genomes with the cores on their source strand. Hits from overlapping chunks that marked the same locus for the same query, species, contig and strand were merged, and distinct paralogous loci were kept. The 20 species comprised 9 non-human primates, 4 non-primate mammals, 3 non-mammalian tetrapods and 4 other vertebrates. Hits with $E \leq 10^{-3}$ and query coverage ≥ 0.8 were high-confidence hits, and the number of species with a high-confidence hit was counted for each lineage group. Cores meeting any of four selection routes were selected (15,913):

- vertebrate_wide: ≥ 1 non-mammalian tetrapod and ≥ 1 other vertebrate
- mammal_conserved: ≥ 3 non-primate mammals and ≥ 3 primates
- primate_focused: ≥ 6 primates and peak P ≤ 0.001
- high_score_rescue: peak P ≤ 0.0005 (no lineage condition)

Candidates meeting several routes were labelled by the first in this order (among the 15,821 candidates with a covariance model (CM; see Alignments): vertebrate_wide 2,110, mammal_conserved 11,140, primate_focused 1,531, high_score_rescue 1,040). The route names are selection strata of the search; they do not imply a lineage-specific origin or absence from other lineages.

**Alignments.** Seed alignments were built from the high-confidence loci of the source-strand search (sequences cut at the Riboseek alignments), and initial covariance models (CMs) were constructed with the human structure as SS_cons and the human columns as RF (`cmbuild --hand`, `cmalign --cyk --notrunc --noprob`).[76] The boundaries of the seed loci were then refined: each high-confidence locus was cropped with one locus length of flank on either side, its boundaries were set by a glocal CYK search with the calibrated initial CM (`cmcalibrate -L 0.01`; `cmsearch --max -g --cyk --acyk --notrunc --toponly -T -1000`), and the sequences were re-aligned to build the seed CM. The 92 selected candidates without a high-confidence locus in the source-strand search (all selected by high_score_rescue; 2 of them had high-confidence hits in the search for the routes) had no seed CM and were not expanded. The remaining 35 genomes of the panel (34 vertebrates and the human genome, searched for paralogous loci) were searched with Riboseek with the source-strand cores, regions of three times the model length around hits with E ≤ 0.1 were cropped, and their boundaries were set in the same way with the seed CM. Hits were accepted with a bit score of at least the 10th percentile of the seed hits minus 10 bits and a model coverage of ≥ 0.8. Added sequences were aligned to the seed with `cmalign --mapali`, yielding alignments of up to 55 species (final55) and their CMs for 15,821 candidates (median 34 species), which form the RLM-Fam candidates. Candidates were grouped into provisional families by reciprocal CM scores; each family has an ID such as RLMFam00092, which is used in the text and in Supplementary Table S4. Candidate pairs were first proposed by an HMM-only search of each seed CM against all human cores (`cmsearch --hmmonly`, E ≤ 10; 177,621 pairs). For each pair of candidates A and B, A's calibrated seed CM scored up to four held-out sequences of B (expansion sequences, one per species, chosen across lineage groups for 4-mer diversity, excluding the seed rows and the human core), and the relative score was $r_{A\to B} = (b_{A\to B} - T_A)/T_A$ ($T_A$, the 10th percentile of seed hits minus 10 bits; $b_{A\to B}$, the median score). Provisional families (14,037) were the connected components of pairs whose relative score averaged over the two directions was at least −0.525; the 21 pairs with no held-out sequences in either direction were left out. This is neither an Rfam gathering threshold nor a P value. Candidates with more than 100 merged high-confidence loci or final55 loci were classed as high-copy.

#### Structural support

**In-cell reactivity.** We used published processed icSHAPE reactivities: in vivo tracks for HEK293, HeLa, K562, HepG2 and H9 from GSE145805, and the HEK293T track from GSE74353.[49,50,78] Reactivities were not recomputed from raw reads. For GSE145805, a core had to lie entirely within a same-strand GENCODE v26 exon and match the transcript sequence exactly. For each candidate and cell line, the isoform with the most measured positions was used. For combinations in which at least 50% of the core was measured and at least 10 paired and 10 unpaired nucleotides were present, the AUROC was computed with unpaired nucleotides as positives and reactivity as the predictor. As a control, the predicted paired/unpaired labels were circularly shifted by every possible offset; the mean AUROC over shifts was the control AUROC, and $(1 + \#\{\text{control} \geq \text{observed}\})/(1 + \text{number of shifts})$ was the empirical P value for each cell line. AUROCs were averaged over cell lines within each core, with cores as the unit of inference (10,000 bootstrap resamples, seed 20260906; paired two-sided Wilcoxon tests between methods). For comparison, the same AUROC was computed on the same cores and measured positions for the RNAfold MFE structure and for structures with only the BPP or only the dependencies as the pair weight. Candidates with an empirical P ≤ 0.05 in at least half of their cell lines were counted as supported by icSHAPE. For GSE74353, candidates were limited to those matching a unique position in Ensembl release 74 transcript sequences. The two data sets can include the same candidates, so we do not call them two independent validation sets.

**Covariation.** Covariation of the proposed base pairs was tested with R-scape 2.6.16 (`-s --nofigures`) under three alignment conditions:[20,51] (i) the final55 alignment made with the CM, (ii) the same sequences realigned with MAFFT and (iii) Riboseek high-confidence hits aligned with MAFFT without structural information.[52] Proposed pairs were mapped to alignment coordinates through the human row. Condition (iii), which depends neither on CM-based sequence selection nor on CM alignment, was the primary condition. Candidates were counted separately at three levels: at least one significant pair, at least two, and at least three with at least two of them in the same helix. Power was expressed as the number of significant pairs expected by R-scape. Structures were never rejected solely because covariation was not significant.

**Covariation background.** Because the call is a test per candidate, the rate of chance positives in the candidate set was estimated with shuffled alignments. Each alignment of conditions (ii) and (iii) was shuffled once with SISSIz 0.1 (`--simulate -d`; phylogeny, gap pattern and dinucleotide content preserved, covariation destroyed), the same SS_cons projected from the human row was put back on the same columns, and R-scape was run with the same settings.[37] Alignments for which SISSIz did not converge (about 8%) were excluded. Candidates and their backgrounds were paired, and intervals for the difference in rates were obtained by bootstraps over candidates (5,000 resamples).

**Structural signal in homologues.** The alignments of condition (iii) (15,809 candidates) were shuffled 19 times each with SISSIz 0.1 (`--simulate -d`).[37] When SISSIz did not converge, it was rerun with its `--flanks` value raised to 48, 96, 192, 384 and 768 in turn; 1,235 candidates (7.8%) for which 19 shuffles could still not be obtained were excluded. The native alignment and its 19 shuffles were scored with RLMHelix-MSA under the settings of Fig. 3 (windows of 200 RF columns moved in steps of 100, pruning to six sequences, 30–300 nt, geometric mean of dependency and RNAalifold BPP, stems of four or more pairs) and with dependencies alone without BPP. Because the human sequence was chosen by the scan score, the main analysis removed the human row before scoring; scores with the human row kept were also computed. An alignment scored above its shuffles when it scored higher than all 19 (empirical P = 1/20), so the chance rate is 5%. The 14,565 candidates for which all 19 shuffles were scored were used (9 were excluded). 95% intervals for the fractions were obtained by bootstraps over candidates (5,000 resamples, seed 20261001). Alignment identity was the mean pairwise identity reported by R-scape. To check the dependence on how the alignments were built, the same scoring was applied to the alignments of condition (i) (built with a covariance model of the predicted structure), using the 14,356 candidates whose 19 shuffles could all be scored with the human row removed.

**Calibration on real alignments.** From the 14,565 candidates, 600 were drawn at random (seed 20261010), each with one matched control of the eCLIP analysis (same gene and strand, same length, region type and exon status, the closest GC content and a low scan score). Both were searched with the same Riboseek settings against the 20 species, the best high-confidence hit per species was aligned with MAFFT (`--auto`), and the alignments were shuffled and scored as above (598 candidates and 596 controls aligned; 593 pairs with all 19 shuffles scored). With the human row removed, 60.0% of the candidates (95% interval 56.0–63.9%) and 20.7% of the controls (17.7–23.9%) scored above all 19 shuffles (63.4% and 16.9% with the human row kept); across alignments of 5 to 20 homologues, 19–22% of the controls scored above all 19 shuffles.

**False discovery rate of the catalogue.** Under SISSIz shuffles, the rank of a native alignment without structure is uniform on 1–20, so Storey's estimate with $\lambda = 0.5$, $\pi_0 = P(\text{rank} \geq 11)/0.5$, estimates the fraction of candidates whose alignments are indistinguishable from their shuffles:[79] 0.133 for the 14,565 candidates (95% interval 0.125–0.141; 2,000 bootstrap resamples of candidates) and 0.135 for the 13,809 families with an alignment (one random member per family, 200 draws); 0.10 for candidates with lfdr ≤ 0.2, 0.07 for those with a BH window, and 0.12, 0.11 and 0.15 for tiers A, B and C. The frequencies of ranks 11 to 20 fall from 1.45% to 0.32% instead of being flat, so the part without signal also ranks above its shuffles more often than uniform ranks allow, as the controls do, and $\lambda = 0.25$ ($P(\text{rank} \geq 6)/0.75$) gives 0.24. With the ranks of the controls as the background, $P_{\text{cand}}(\text{rank} \geq 11)/P_{\text{ctrl}}(\text{rank} \geq 11)$ is 0.25 (0.22–0.30). The false discovery rate of the catalogue therefore lies between about 0.13 and 0.25, depending on the background. The controls had slightly fewer homologues than the candidates (mean 10.1 versus 11.8 species), but the fraction of controls scoring above all 19 shuffles did not change with the number of homologues (19–22% for 5–20 homologues), so this difference has little effect on the estimate.

#### Comparison with earlier genome screens

**Prediction sets.**

- RNAz 2005: the authors' hg17 BED files (Set1–3, P > 0.5 and P > 0.9) were lifted to hg38 with UCSC liftOver (minMatch 0.95).[13,88] The input regions of the original screen (the 2005 PhastCons most-conserved elements) are no longer distributed, so we approximated them with hg17 phastConsElements17way elements of at least 50 nt lifted to hg38 (113 Mb; the original input was about 83 Mb).[48] Because the approximation is larger, the fraction of candidates outside the input is a conservative estimate.
- EvoFold: the original predictions (UCSC hg19 `evofold`, 47,510) and the genome-wide v2 predictions (hg18 `evofoldV2`, 37,381) were each lifted to hg38 (more than 99.9% converted).[11,14]
- CMfinder CRSs (Seemann 2017): we used Supplementary Data 1 of release v2.1, with the 2020 correction of hg38 coordinates, downloaded from the authors' website. Two sets were taken from it: all published CRSs (RNAPhylo score > 50; 773,375 CRSs in 515,335 regions on primary chromosomes) and the 156,171 with a reported FDR of at most 10%.[15]

**Overlaps and background.** Because RNAz 2005 predictions are unstranded, the primary analysis counted overlaps of at least 1 bp on either strand; same-strand overlaps with EvoFold and CRSs were counted as a sensitivity analysis.[77] The background was the overlap fraction of intervals with the same lengths as the candidates or BH loci, placed at random within the scanned space (the merged eligible windows) on the same chromosome (`bedtools shuffle -incl -chrom`; 20 replicates for candidates and 100 for BH loci). Enrichment by window rank was computed on the 6,924,574 windows collapsed per coordinate to the P value of the better strand, split into the top 0.1%, the top 0.1–1% and deciles of the rest, as fold over all windows. Candidates overlapping none of the four prediction sets were split into those within the approximate RNAz input and those outside it (groups as in Fig. 4h), and their icSHAPE agreement and covariation under condition (iii) were compared with those of candidates overlapping a prediction (Mann–Whitney tests for icSHAPE agreement, Fig. 5a; Fisher's exact tests for covariation, Supplementary Fig. S7c).

**Rfam enrichment along the selection**

For every selection stage (all eligible windows, top-1% windows, cores, selected cores, RLM-Fam candidates, and BH windows and loci), the fraction of intervals overlapping Rfam was counted under one definition. Rfam was the set of significant Rfam 15.1 full hits on GRCh38 primary chromosomes (all families and all lengths; 20,516 hits in 1,077 families).[41] The primary rule was an overlap of at least 1 bp on the same strand, with either-strand overlap as a sensitivity analysis. Three backgrounds were used: (a) fold and odds ratio relative to all eligible windows; (b) the stage's intervals placed at random within the scanned space with the same lengths, strands and chromosomes (`bedtools shuffle -incl`, with the complement given to `-excl` so that every placed interval lies within the scanned space; 20–1,000 replicates); and (c) the intervals placed within random eligible windows of the same chromosome and strand (a sensitivity analysis that preserves the window distribution). Intervals for fractions are Wilson 95% intervals. Sensitivity analyses are shown in Supplementary Fig. S5.

**Comparison with thermodynamic scanning on the scan windows (Fig. 4e, f)**

Of the 7,869,821 eligible scan windows, those overlapping a significant Rfam 15.1 full hit on the same strand by at least 1 bp (4,575) were positives, and all windows were scored. For each window sequence (in the scan's orientation) we computed the RNAfold MFE (ViennaRNA 2.7, default parameters) and the ScanFold 2.0 z-scores as above (mononucleotide- and dinucleotide-shuffle models; 200 nt lies within their training range). Rankings by each score gave the fraction of positives among the top 0.1% and 1% of windows, the enrichment over all windows and the AP. To count each known RNA once, for the Rfam hits at least half covered by eligible windows (1,463) we computed the fraction whose best window ranked in the top 1%, overall and by RNA class. Intervals are from a block bootstrap over 1-Mb genomic blocks (200 replicates, seed 20261003).

**Comparison with RNA-binding protein binding (eCLIP)**

**Data.** We used reproducible peaks (IDR, GRCh38) from ENCODE eCLIP in K562 and HepG2 (250 experiments, 168 RBPs).[54,80] All overlaps were counted on the same strand, except in the comparison in the format of Seemann 2017 (below), which used interval midpoints on either strand.

**Matched within-gene controls.** Each candidate core was matched with up to five control intervals that:

- lie on the same chromosome and strand, with the centre of their scan window inside the core's representative gene (GENCODE v50);
- have the same region type (intron, lncRNA, 3′UTR and so on) and the same exon overlap status;
- have an RLMHelix score below the genome-wide median and a calibrated P > 0.1;
- overlap no candidate core on either strand;
- are cut from the window centre with the core's length and, among the qualifying windows, have the GC content closest to the core's.

For each experiment, a Mantel–Haenszel odds ratio and a Cochran–Mantel–Haenszel test were computed with each candidate and its controls as one stratum, and P values were BH-adjusted across experiments (Supplementary Table S5).[81] For comparison, we also ran an analysis in the format of Seemann 2017 (interval midpoints, either strand, binomial tests within GC bins) with all scan windows as background.[15]

**Dose–response and dsRNA-binding proteins.** At the window level, Mantel–Haenszel odds ratios were computed for bins of calibrated P value, stratified by gene × region type × exon overlap × GC. RBPs that recognize double-stranded RNA (DGCR8, DROSHA, ILF3, STAU2, DHX30 and XPO5) were defined as a group before per-RBP results were examined. An analysis excluding candidates within 500 nt of a GENCODE miRNA gene was also performed.

**Genome-browser views (Fig. 6c, Supplementary Fig. S11).** For ENCODE eCLIP in HepG2 (in K562 for Supplementary Fig. S11d), the strand-specific signal of unique reads (bigWig) of the two IP replicates and the size-matched input, together with the IDR peaks, were drawn on the coordinates of each candidate with the MANE Select gene structure and the calibrated P values of windows in the top 5%. The IP signal is the mean of the two replicates. In the zoom, RLMHelix base pairs are drawn as arcs and the signal of the first RBP named for each candidate in the legends (for example, STAU2 for *NUTF2*) is drawn downwards. The tracks show read coverage, not crosslink positions. Candidates on the minus strand are drawn with reversed coordinates, so that every view runs 5′→3′ from left to right. For *HMGCS1*, cleavage sites from PolyASite 2.0 (GRCh38, Ensembl 96)[57] carrying at least 1% of the expression in view are drawn, and the zoom marks UGUA and AAUAAA (AUUAAA) motifs. The positive control was DGCR8 eCLIP at the candidate that overlaps the pri-miR-22 hairpin. Accessions of the files used are listed in Supplementary Table S8.

#### Per-candidate binding partners and the chance ratio

For each pair of candidate and eCLIP experiment, three calls were made:

- **bound:** a peak overlaps the candidate core;
- **control-free:** in addition, none of the matched within-gene controls (at least two) is bound;
- **candidate-specific:** in addition, the candidate's peak is the strongest of that experiment within the host gene (log2 fold enrichment over input).

Candidates whose host gene is not expressed in the cell line (mean TPM < 1 over six replicates of ENCODE non-target control RNA-seq (four shRNA and two CRISPR controls per cell line)) were excluded as not measurable rather than counted as unbound. For each RBP, a rarity weight $\ln[(N_{\text{tested}} + 1)/(N_{\text{specific}} + 1)]$ was computed, where $N_{\text{tested}}$ is the number of candidates measurable for that RBP and $N_{\text{specific}}$ the number with candidate-specific binding.

To estimate how many calls arise by chance, for candidates with at least three controls the first control served as a pseudo-candidate, and the same calls were made with the remaining controls (6,088 pairs). Because candidates and pseudo-candidates are compared within the same gene and experiment, differences between them reflect candidate status rather than the gene or the experiment. For each RBP, discordant calls between candidates and pseudo-candidates were compared with McNemar's test (exact binomial) and BH-adjusted across RBPs.[82]

#### Recovery of known functional structures

We collected cis-acting structures in human mRNAs and lncRNAs whose function and binding partner have been established in the literature (12 classes, 82 elements, 73 genes), including *MAT2A* hairpins and METTL16, histone stem-loops and SLBP, IREs and IRPs, and SECIS elements and SECISBP2. The 31 papers supporting these elements were all checked against their PubMed records, including titles (Supplementary Table S3). Coordinates were set mainly from Rfam 15.1 hits within the genes in which function had been shown, and otherwise from the literature or by sequence match (the method is recorded for each element). For each element, we determined whether same-strand eligible windows covered at least 50% of it, the rank of the calibrated P value of its best window among all windows, and whether it fell in the top 1% and in RLM-Fam. The background was 100 intervals of the same length placed within the same gene (matching exonic status and at least 200 nt from the element), or within 20 kb on either side of the element when the gene was too short. Counts in the top 1% and in RLM-Fam were compared with the mean background rate by binomial tests, and ranks with the median of each element's background by paired Wilcoxon tests. Whether the known partner could be named was assessed from the per-candidate binding calls when the partner had eCLIP data, and otherwise from the literature-defined structure–sequence rules (Methods). Enrichment at thresholds from the top 0.01% to 50% was computed on the same scale for Rfam hits (300 miRNA precursors, 295 C/D snoRNAs, 189 H/ACA snoRNAs, 66 tRNAs and 122 cis-regulatory elements; up to 300 per class sampled from hits inside the scanned space) (Supplementary Fig. S6).

#### Consistency with known partner properties

We counted the candidate-specific calls that agree with a known property of the partner. For sequence-specific RBPs, recognition motifs were the ten 5-mers with the highest enrichment among those with $z \geq 3$ at the most enriched protein concentration in the 5-mer enrichment tables of ENCODE RNA Bind-n-Seq (RBNS; 96 RBPs).[89] A call was consistent when the interval carried a motif with at least one occurrence whose five nucleotides were unpaired in both the RLMHelix and MFE structures (21 of the 23 RBPs shared with eCLIP; FUS has no 5-mer with $z \geq 3$, and RBM22 was counted as a spliceosomal factor). For double-stranded RNA-binding proteins (dsRBPs), consistency required a helix of at least 12 directly stacked pairs in the MFE structure of the interval; for the 12 spliceosomal factors, the interval had to span, or lie within 100 nt of, a same-strand internal exon boundary. Pseudo-candidate structures were recomputed with the student used in the genome scan, in the context of their scan windows; applied to 60 candidates, the same recomputation reproduced their catalogue structures exactly. For consistent calls, the pseudo-candidate rate divided by the candidate rate was taken as the chance ratio (bootstrap intervals, Supplementary Fig. S10). To see how far the property itself differs between candidates and pseudo-candidates, we compared the fraction of measured candidates and of measured pseudo-candidates that meet it (the dsRBP property: 28.7% of candidates, 1.9% of pseudo-candidates). For dsRBPs, the fraction of candidate-specific calls that meet the property was tested with a one-sided binomial test against its fraction among unbound measured candidates (DGCR8: 5 of 7, expected 28.7%).

#### Implication table and evidence tiers

**Implication table.** From these analyses, we selected candidates with grounds for a pairing of structure and partner and recorded the literature support at five levels (Supplementary Table S6):

- reported: the element, or its regulation by the partner, has been reported;
- reported for the gene: regulation of the host gene by this mechanism has been reported, but the role of this structure has not;
- analogous precedent: the same partner has been reported to act on a similar structure in another gene;
- consistent with partner properties: the pairing agrees with known properties of the partner, but the element has not been reported;
- partner unidentified: a conserved structure of a type that a partner could recognize, without binding data.

All PMIDs were checked on PubMed.

**Evidence tiers.** Each catalogue candidate was annotated with statistical support (lfdr $\leq$ 0.2, which all candidates containing a BH-accepted window meet) and independent structural support (icSHAPE support or covariation under condition (iii)). Candidates with both were assigned to tier A, those with one to tier B and those with neither to tier C (Supplementary Table S4). Absence of covariation and missing icSHAPE measurements were not treated as evidence against a candidate. Rfam overlap is listed only as a descriptive column and was not used to define tiers.

**Pull-down mass spectrometry (Fig. 6d–f, Supplementary Figs S12 and S13)**

**Baits.** The 24 baits, chosen independently of this scan, were 16 EvoFam elements, including known RNAs such as let-7, the *MAT2A* hairpins and SNORA2,[14] and 8 conserved structures found with CMfinder by Hou et al. (hereafter CM-line; 6 human, 2 mouse)[65] (Supplementary Table S7). Bait RNAs were transcribed in vitro with the MEGAshortscript T7 Transcription Kit (Thermo Fisher Scientific).

**Preparation of FLAG-conjugated bait RNA.** The transcribed RNAs were conjugated with FLAG at the 3′ end as described previously,[90] with minor modifications. Briefly, 50 pmol of purified RNA was incubated with 12 $\mu$L of 0.1 M sodium periodate ($NaIO_4$) at 0 °C for 10 min to oxidize the 3′-terminal ribose. The oxidized RNA was precipitated with ice-cold 2% $LiClO_4$/acetone and washed with 1 mL of ice-cold acetone. After brief air-drying, the RNA pellet was dissolved in 2 $\mu$L of 0.1 M sodium acetate buffer, pH 5.2, and mixed with 2 $\mu$L of 30 mM FLAG-hydrazide. The reaction mixture was incubated at room temperature for 30 min and then reduced by adding 2.4 $\mu$L of 1 M sodium cyanoborohydride and incubating for a further 30 min at room temperature. The FLAG-conjugated RNA was purified with RNAClean XP (Beckman Coulter) and used for RNA pull-down.

**RNA pull-down.** 293T cells were lysed in lysis buffer containing 10 mM HEPES, pH 7.5, 150 mM NaCl, 50 mM NaF, 1 mM $Na_3VO_4$, 5 $\mu$g/mL leupeptin, 5 $\mu$g/mL aprotinin, 3 $\mu$g/mL pepstatin A, 1 mM PMSF and 1 mg/mL digitonin. Lysates were cleared by centrifugation at 4 °C, and the supernatants were incubated with FLAG-conjugated bait RNA, anti-FLAG M2 antibody and Protein G magnetic beads for 1 h at 4 °C with gentle rotation. The beads were then collected and washed three times with wash buffer containing 10 mM HEPES, pH 7.5, 150 mM NaCl and 0.1% Triton X-100. Bound RNA–protein complexes were eluted with FLAG elution buffer containing 0.5 mg/mL FLAG peptide, 10 mM HEPES, pH 7.5, 150 mM NaCl and 0.05% Triton X-100. The eluted proteins were processed by the SP3 method:[83] proteins were reduced with TCEP, alkylated with iodoacetamide, captured on paramagnetic beads under high-organic-solvent conditions and digested with lysyl endopeptidase and trypsin. The resulting peptides were recovered, desalted and used for LC–MS/MS.

**LC–MS/MS.** Peptides were analysed on an Evosep One liquid chromatography system coupled to an Orbitrap Astral mass spectrometer (Thermo Fisher Scientific). Desalted peptides were loaded onto Evotips according to the manufacturer's protocol and separated with the standard 100-samples-per-day method on an Evosep-compatible C18 analytical column, with 0.1% formic acid in water as solvent A and 0.1% formic acid in acetonitrile as solvent B. Eluted peptides were introduced directly into the mass spectrometer through a nano-electrospray ion source, and data were acquired in data-independent acquisition (DIA) mode: MS1 spectra in the Orbitrap analyser and DIA MS/MS spectra in the Astral analyser, covering a precursor range of approximately m/z 380–1000 with 2-Th isolation windows. Each bait was pulled down twice, and each pull-down was measured twice (four runs per bait, 96 runs in total).

**Protein identification and quantification.** Raw data were processed with DIA-NN (version 2.0.2).[84] Spectral library generation and database search were performed against the UniProt human reference proteome (UP000005640; release 2021_03, including isoforms) with trypsin as the enzyme and up to two missed cleavages; carbamidomethylation of cysteine was a fixed modification, and oxidation of methionine and protein N-terminal acetylation were variable modifications. Precursor, peptide and protein identifications were controlled at a false discovery rate of 1%. Protein quantities were calculated with the MaxLFQ algorithm[85] implemented in DIA-NN, and the protein-level MaxLFQ intensity matrix was used for the analyses below.

**Binder calls.** MaxLFQ intensities were log2-transformed and median-normalized per sample, a proDA model with a mean per bait and an effect of the pull-down replicate was fitted to the 96 runs,[86] and each bait was compared with the mean of the other 23 by a Wald test. Proteins with adjusted $P < 0.05$ and log2 fold change $\geq 2$ were taken as binders of that bait.

**Baits and labels for the score comparison.** The comparison with the scores used the 16 human baits of at least 30 nt for which alignments could be built (10 EvoFam, 6 CM-line). The 2 mouse baits were excluded because their alignments are the CMfinder alignments of the human baits with another reference row, and the 6 baits shorter than 30 nt lie outside the range of the student and of RLMHelix-MSA (30–300 nt). Double-stranded RNA-binding proteins were ADAR, ADARB1, ADARB2, DGCR8, DHX9, DHX30, DICER1, DROSHA, EIF2AK2, ILF3, PRKRA, STAU1, STAU2, STRBP, TARBP2, ZFR and XPO5 (the six used in the eCLIP analysis are all on this list), and their number among the binders of each bait was counted for Supplementary Fig. S13a. Proteins with a Pfam double-stranded RNA-binding domain were also counted for the labels and for Supplementary Figs. S12 and S13b, e (Methods). Known partners were LIN28A and LIN28B for let-7, METTL16 for the *MAT2A* hairpins, DKC1, NOP10, NHP2 and GAR1 for SNORA2, DGCR8 and DROSHA for miR-9-2, and SLBP for the histone stem-loop.

**Scoring.** EvoFam baits used multiz100way alignments projected onto human coordinates, as in Fig. 3; CM-line baits used the original CMfinder alignments of Hou et al., with the human row identical to the bait as reference. Each alignment was compared with 100 SISSIz shuffles (`--simulate -d`)[37] and scored with RLMHelix-MSA, dependency only, RNAalifold BPP only, RNAalifold −MFE and RNAz with the settings of Fig. 3. Single sequences were scored with the four scores of Fig. 2 (RLMHelix, dependency only, BPP only and RNAfold −MFE) and compared with 100 uShuffle shuffles ($k = 2$).[36] As supplementary analyses, single-sequence RLMHelix was also computed for the 2 mouse baits (their rows in the alignments of Hou et al.) with the student and for the 6 baits shorter than 30 nt with the teacher.

**Comparison with the scores.** To compare baits, each score was converted to z = (score − mean of shuffles) / s.d. of the shuffles of that bait. We computed ROC curves and the AUC for baits with versus without a structure-recognizing binder (at least one double-stranded RNA-binding protein, which reads duplexes, or one of the known partners listed above; two-sided Mann–Whitney test) and the Spearman correlation of z with the number of double-stranded RNA-binding proteins (also as a partial correlation adjusted for bait length), with 95% intervals from 10,000 bootstrap resamples of baits (seed 20261002). Differences in AUC between scores on the same baits were tested with the paired DeLong test.[87] As a comparison without a threshold, the Spearman correlation of z with the mean log2 fold change against the other 23 baits over all quantified double-stranded RNA-binding proteins (the double-stranded RNA-binding enrichment) was also computed. As a control, the same quantity was computed over all quantified RNA-binding proteins (UniProt RNA-binding keywords), to ask whether the scores follow RNA binding in general or binding by proteins that recognize structure. Typical skin and secretion contaminants (61 proteins defined by gene-symbol rules: keratins, desmosome and cornified-envelope proteins, S100A7/8/9, SERPINB3/4, DCD, LCN1 and others) were kept in the data, the binder counts and the labels, and were only drawn with a separate symbol in the volcano plots. For the structure-recognizing binder, proteins with a Pfam double-stranded RNA-binding domain in UniProt (dsrm, DZF, Staufen_C, DND1_DSRM and zf-met) were also counted as double-stranded RNA-binding proteins (DZF occurs in double-stranded RNA-binding proteins such as ILF3 and ZFR; zf-met is the matrin-type zinc finger with which JAZ, also called ZNF346, binds double-stranded RNA). As a sensitivity analysis, proDA was refitted to the 64 runs of the 16 baits, and binders were called by comparing each bait with the other 15; as a check of this refitting, applying it to all 24 baits reproduced the original labels of all 16 baits.

#### Base-pair prediction on bpRNA (Supplementary Fig. S1)

We used the 5,313 sequences of 30–300 nt among the 5,401 sequences of bpRNA-new.[35] For the teacher and the student, the geometric mean of APC and BPP was decoded with the non-crossing DP and isolated pairs were removed; the RNAfold control used the MFE structure. Predicted and reference pairs were matched exactly, and per-sequence F1 scores were averaged. Differences between methods were estimated by bootstraps over sequences (10,000 resamples, seed 20260909). These sequences may be part of the language model's pretraining data, so the comparison tests whether distillation preserves base-pair information, not generalization to unseen RNAs.

**Software**

- Structure and alignment: ViennaRNA 2.7, RNAz 2.1, SISSIz 0.1, uShuffle 1.0, Infernal 1.1.5, MAFFT 7.526, R-scape 2.6.16, R2R (distributed with R-scape 2.6.16)
- Phylogenetic models: CMfinder 0.4.1.18 (RNAPhylo), EvoFoldV2 (phy library, commit `10ee240d`)
- Sequence search: Riboseek (commit `eba085cb`)
- Genomic intervals: bedtools 2.31.1, UCSC liftOver
- Repeat filtering: tantan 51, Inverted Repeats Finder 3.09
- Language model: PyTorch 2.11, transformers 5.5, multimolecule 0.0.9 (RiNALMo-giga)
- Composition-calibrated MFE: ScanFold 2.0 (published models, commit `5eec7c20`)
- Statistics: SciPy 1.17.1, scikit-learn 1.8.0
- Mass spectrometry: DIA-NN 2.0.2, proDA 1.12.0

### Supplementary figures

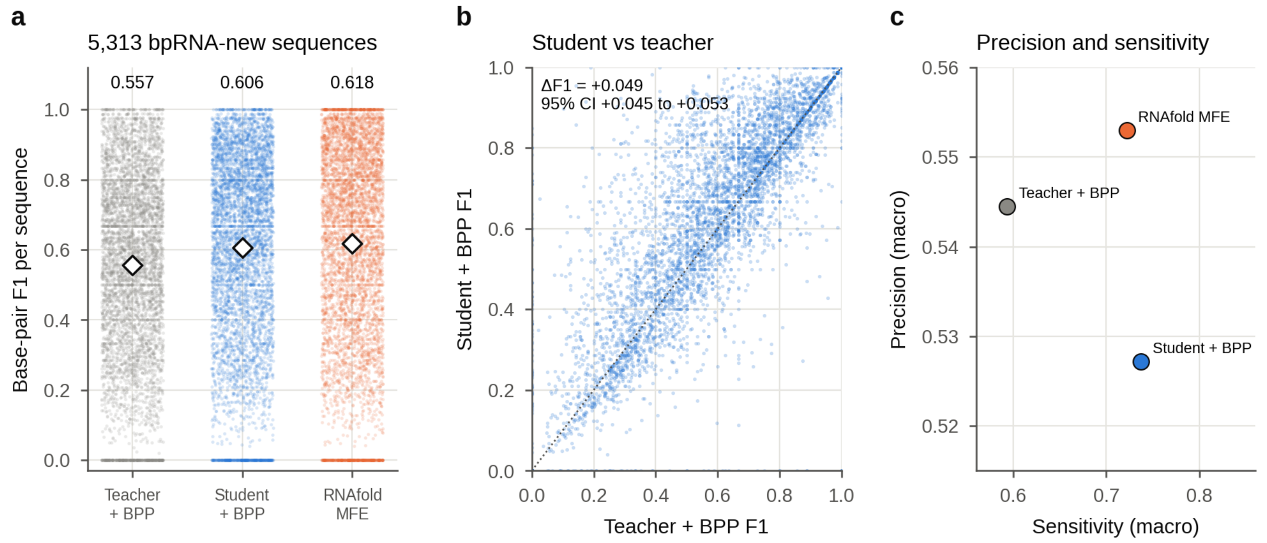


**Supplementary Fig. S1 | The distilled student retains base-pair information.** Base-pair prediction for 5,313 bpRNA-new sequences (30–300 nt). The geometric mean of the APC-corrected dependency and the RNAfold BPP was decoded by non-crossing Nussinov-type DP, and isolated base pairs were removed. **a**, F1 per sequence and means (diamonds): teacher 0.557, student 0.606, RNAfold MFE 0.618; student minus RNAfold MFE, −0.012 (95% interval −0.016 to −0.008; paired bootstrap as in **b**). **b**, Student versus teacher per sequence; mean difference +0.049 (95% interval 0.045–0.053; 10,000 paired bootstrap resamples of sequences). **c**, Precision and sensitivity averaged over sequences.

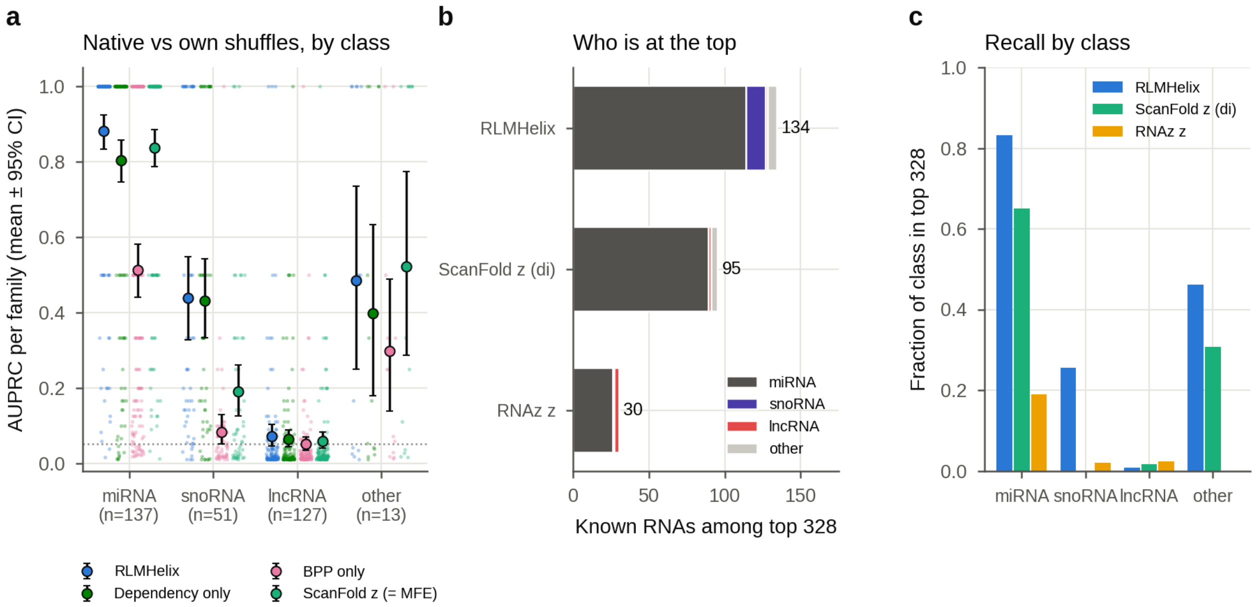


**Supplementary Fig. S2 | RLMHelix gains most for snoRNAs, and lncRNA families are barely distinguished by any score.** The 328 Rfam families split by Rfam family type (miRNA 137, snoRNA 51, lncRNA 127, other 13). **a**, AUPRC of each family (small points) and their mean with 95% intervals from 2,000 bootstrap resamples of families (circles and lines) by class; for snoRNAs, RLMHelix 0.438 and ScanFold 2.0 z 0.189. Families annotated as 'lncRNA' in Rfam are barely distinguished by any score (AUPRC 0.072 or less; dotted line, chance expectation 0.051). **b**, Composition of the known RNAs among the top 328 sequences (as many as there are known RNAs) in a single ranking of all sequences: 134 for RLMHelix (miRNA 114, snoRNA 13), 95 for ScanFold 2.0 z (miRNA 89, snoRNA 0) and 30 for RNAz z. **c**, Fraction of the known RNAs of each class that rank among the top 328 (recall).

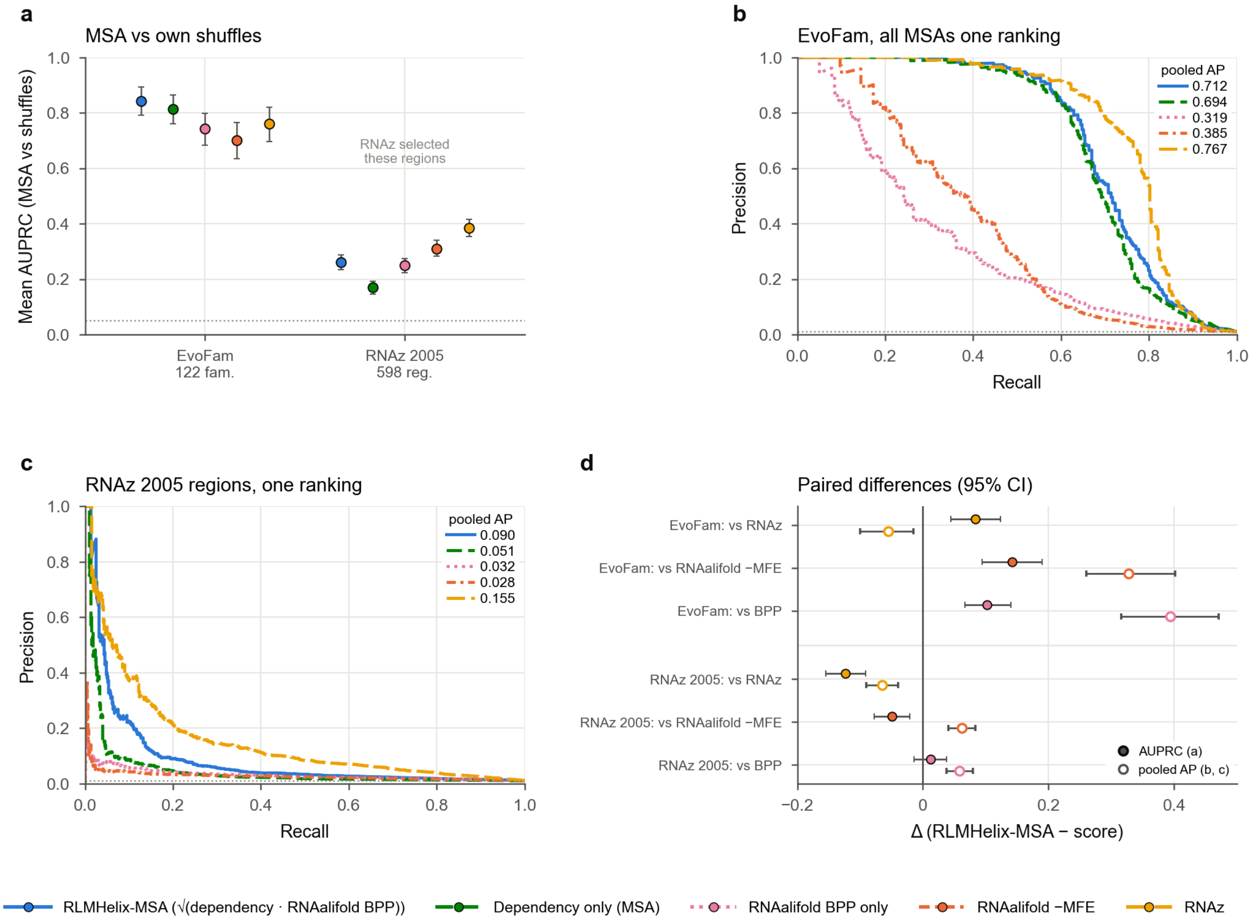


**Supplementary Fig. S3 | Agreement of RLMHelix-MSA with the predictions of earlier screens.** EvoFam (GW set; 318 loci in 122 families) and 598 regions drawn by stratified sampling from the RNAz screen of the human genome (RNAz 2005) are predictions of earlier screens, not sets of true structures, so only agreement can be compared; the RNAz 2005 regions were selected by RNAz itself, so the comparison with RNAz is biased in its favour. The phylogenetic models (RNAPhylo, EvoFold) were scored on Rfam only (Fig. 3). Each MSA was compared with 100 SISSIz shuffles, as in Fig. 3. **a**, Mean AUPRC of MSAs against their own shuffles, with 95% intervals (dotted line, chance expectation 0.051). For EvoFam and RNAz 2005, respectively: RLMHelix-MSA 0.843 and 0.261; dependency only 0.813 and 0.169; BPP only 0.742 and 0.249; RNAalifold −MFE 0.702 and 0.310; RNAz 0.760 and 0.384. **b**, **c**, Precision–recall curves for all MSAs of EvoFam (**b**) and RNAz 2005 (**c**) in a single ranking; the legend gives the pooled AP of each curve, with line styles as in the key under the figure (EvoFam: 0.712, 0.694, 0.319, 0.385, 0.767; RNAz 2005: 0.090, 0.051, 0.032, 0.028, 0.155). **d**, Paired differences between RLMHelix-MSA and the other scores with 95% intervals (filled, mean AUPRC of **a**; open, pooled AP of **b** and **c**). Against RNAz: +0.083 (0.044–0.124) and −0.055 (−0.100 to −0.016) for EvoFam, and −0.123 (−0.155 to −0.091) and −0.065 (−0.090 to −0.040) for RNAz 2005. Against RNAalifold −MFE (mean AUPRC): +0.142 (0.094–0.190) for EvoFam and −0.049 (−0.077 to −0.021) for RNAz 2005. Intervals are 95% bootstrap intervals (10,000 resamples for mean AUPRC and 2,000 for pooled AP; units, families for EvoFam and regions for RNAz 2005).

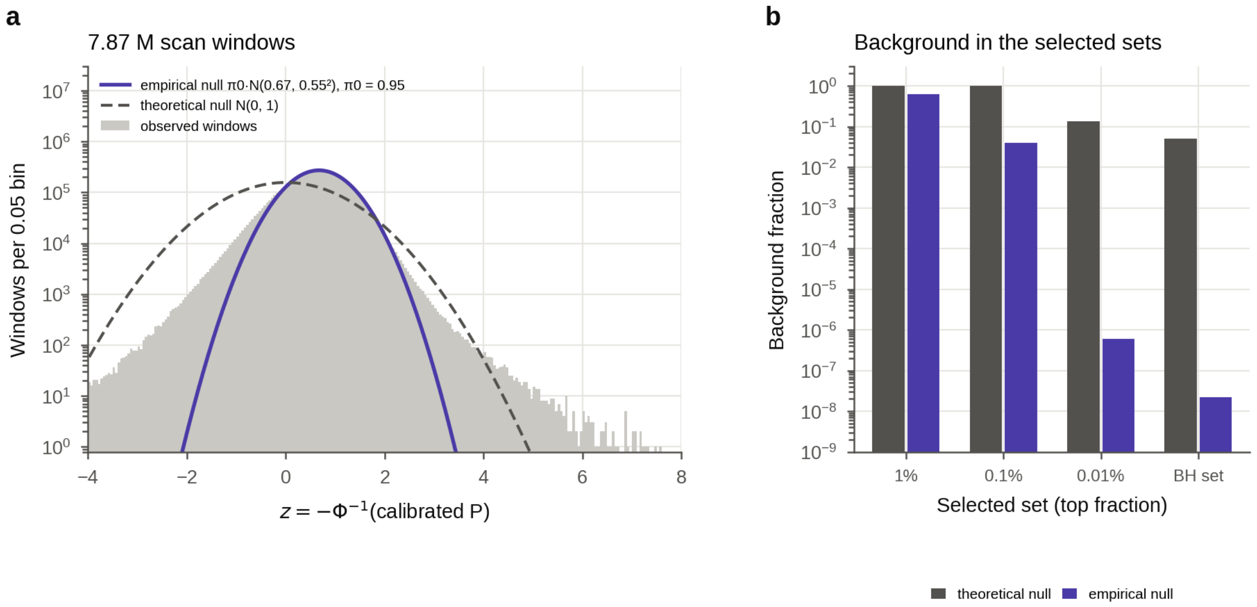


**Supplementary Fig. S4 | Empirical null of the genome scan.** **a**, Distribution of $z = -\Phi^{-1}(p)$ from the calibrated P values of the 7,869,821 windows (grey, log scale), the empirical null $\pi_0 \cdot N(0.673, 0.548^2)$ fitted to the central part (z 0.22–1.01; $\pi_0$ = 0.951; violet) and the theoretical null N(0, 1) (dashed). **b**, Estimated fraction of background windows in the selected sets (as in Fig. 4c). Under the theoretical null, 1.0 for the top 1% and 0.1%, 0.13 for the top 0.01% and 0.050 for the BH set; under the empirical null, 0.61, 0.039, $6 \times 10^{-7}$ and $2 \times 10^{-8}$, respectively. Empirical estimates beyond the top 0.1% depend on extrapolating the normal tail and are uncertain; 0.050 for the BH set is the nominal value.

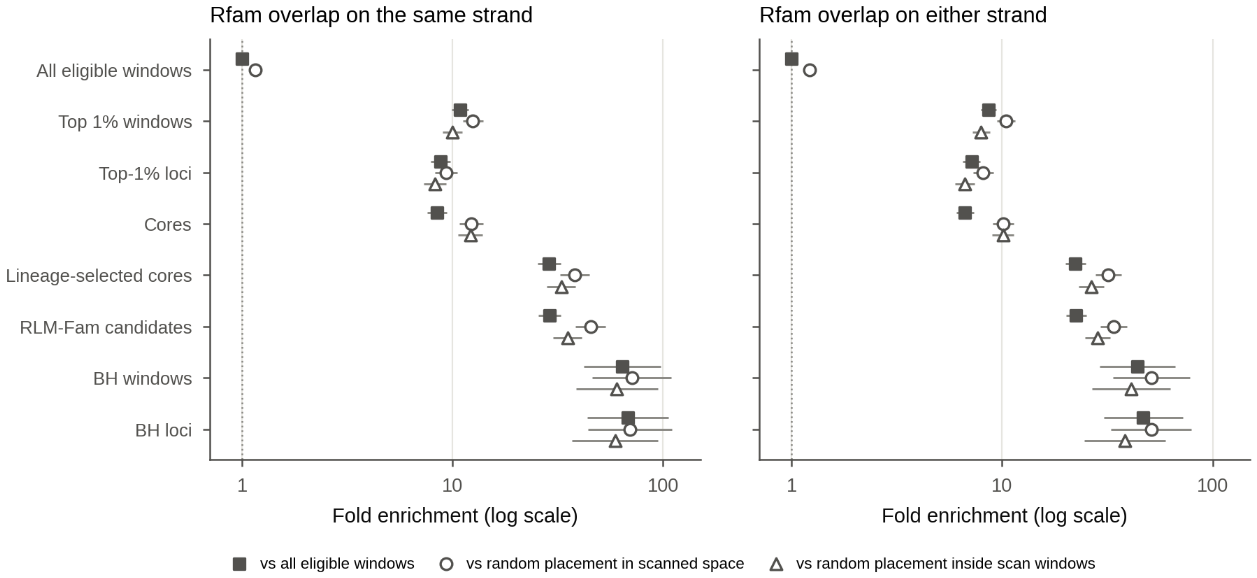


**Supplementary Fig. S5 | The stage-wise Rfam enrichment holds against three backgrounds.** The enrichment of Fig. 4d against three backgrounds: fold over all eligible windows (filled squares), over intervals of the same lengths placed at random in the scanned space (open circles) and over intervals placed at random inside scan windows (triangles). Lines, 95% intervals (Katz log intervals of the ratio of fractions). Random placements: 20 for the top 1% of windows, the top-1% loci and the cores, 50 for the lineage-selected cores and the candidates, and 1,000 (circles) or 500 (triangles) for the BH windows and loci. Sets (n): all eligible windows (7,869,821), top 1% of windows (78,699), top-1% loci (71,459), cores (71,196), lineage-selected cores (15,913), RLM-Fam candidates (15,821), BH windows (565) and BH loci (481). Left, same-strand overlap; right, overlap on either strand. On the same strand, the top 1% of windows is enriched 11-, 12- and 10-fold, RLM-Fam candidates 29-, 45- and 35-fold and the 481 BH loci 68-, 70- and 60-fold.

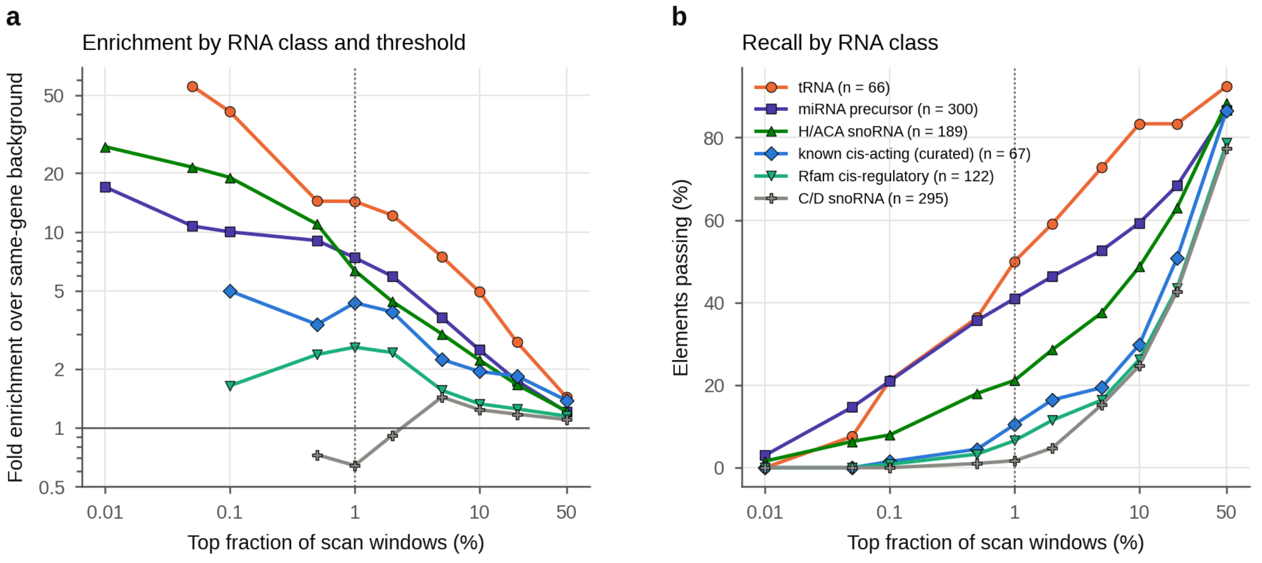


**Supplementary Fig. S6 | Enrichment by the scan differs strongly between RNA classes.** The rank of the best window of each element was compared with 100 intervals of the same length placed in the same host gene (intervals drawn anew, so the expectations differ slightly from those given in the text). Elements: the 67 known cis-acting structures in the scanned space and Rfam 15.1 hits in the scanned space (tRNA 66, miRNA precursors 300, H/ACA snoRNAs 189, C/D snoRNAs 295, cis-regulatory 122; up to 300 per class). **a**, Fold enrichment over the background expectation by threshold. In the top 1%: tRNAs 14-fold, miRNA precursors 7.4-fold, H/ACA snoRNAs 6.4-fold, known cis-acting structures 4.3-fold, Rfam cis-regulatory elements 2.6-fold and C/D snoRNAs 0.6-fold. In the top 0.1%: tRNAs 41-fold, H/ACA snoRNAs 19-fold and miRNA precursors 10-fold. **b**, Fraction of elements passing each threshold. In the top 1%: tRNAs 50%, miRNA precursors 41%, H/ACA snoRNAs 21%, known cis-acting structures 10%, Rfam cis-regulatory elements 7% and C/D snoRNAs 2%. The scan is strong for structures with several stable stems or long helices and weak for short cis-regulatory hairpins and C/D snoRNAs.

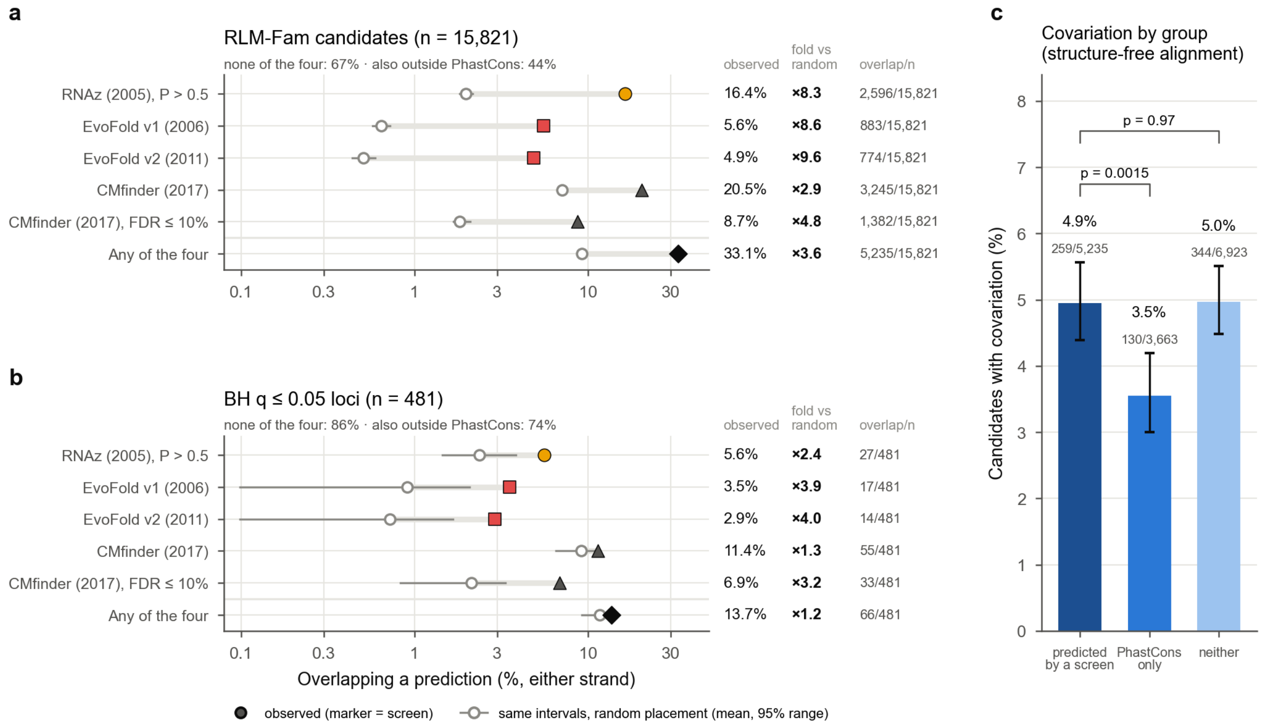


**Supplementary Fig. S7 | Most candidates and BH loci lie outside the predictions of the four earlier screens. a**, Fraction of the 15,821 RLM-Fam candidates overlapping each prediction set (filled; either strand, ≥1 nt) and of intervals of the same lengths placed at random in the scanned space (open; mean and 2.5–97.5 percentiles of 20 placements). Numbers at right give the observed fraction, the fold over random placement and the overlapping candidates: RNAz 2005 16.4% (8.3-fold), EvoFold v1 5.6% (8.6-fold), EvoFold v2 4.9% (9.6-fold), all CMfinder CRSs 20.5% (2.9-fold), CRSs with FDR ≤ 10% 8.7% (4.8-fold) and any of them 33.1% (3.6-fold). Of the candidates, 67% overlap none of the four, and 44% are also outside the PhastCons conserved elements (approximating the input of RNAz 2005 and EvoFold). **b**, The same for the 481 BH-accepted loci (100 random placements): RNAz 2005 5.6% (2.4-fold), EvoFold v1 3.5% (3.9-fold), v2 2.9% (4.0-fold), all CRSs 11.4% (1.3-fold), CRSs with FDR ≤ 10% 6.9% (3.2-fold) and any of them 13.7% (1.2-fold); all CRSs and any of them lie within the 95% range of random placement. Of the loci, 86% overlap none of the four, and 74% are also outside PhastCons. **c**, Fraction of candidates with structure-free covariation, by relation to the four screens (Wilson 95% intervals): overlapping a prediction 4.9% (n = 5,235), in PhastCons without a prediction 3.5% (3,663), neither 5.0% (6,923). P values are from two-sided Fisher's exact tests against the candidates overlapping a prediction and are not adjusted for multiple comparisons. Restricted to the same strand, candidates overlap EvoFold v1 in 3.4%, v2 in 3.1%, all CRSs in 13.5% and CRSs with FDR ≤ 10% in 5.6%.

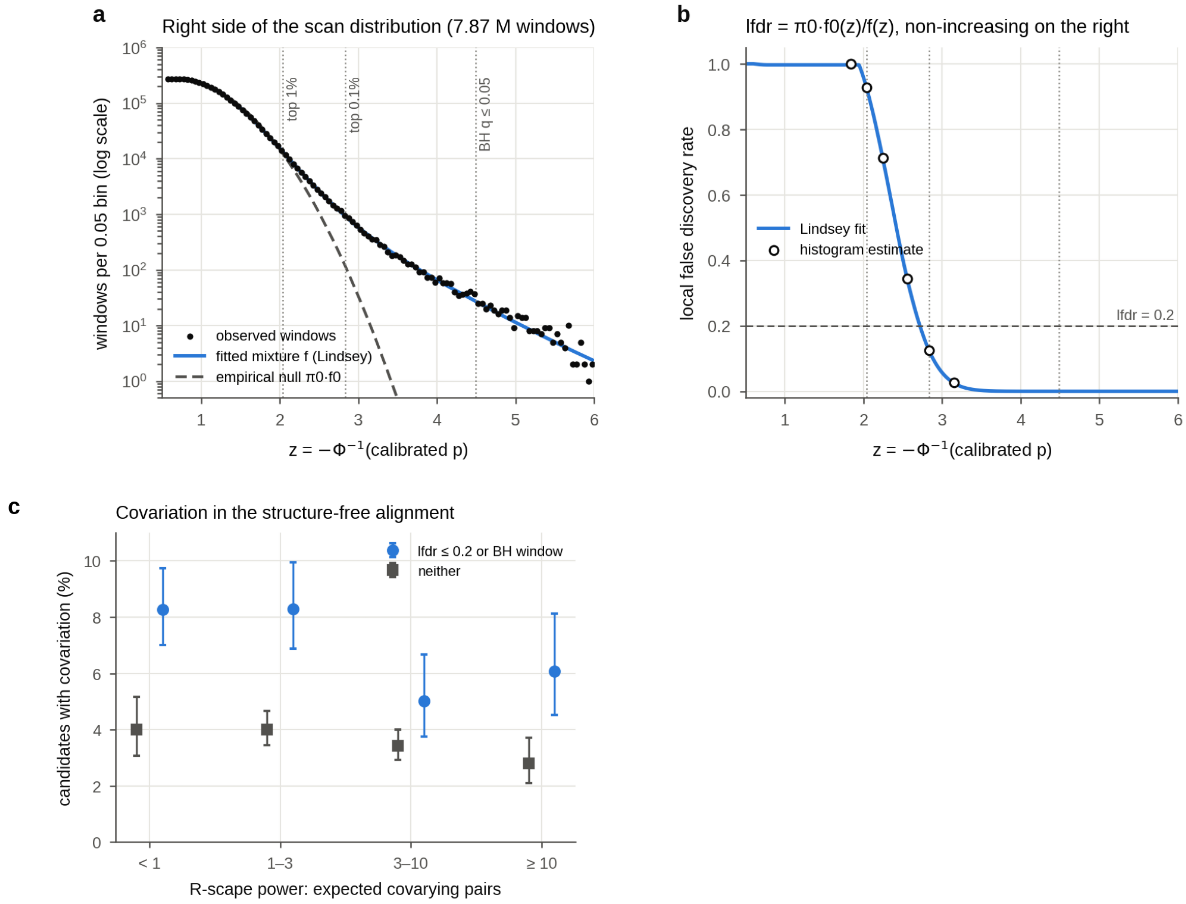


**Supplementary Fig. S8 | Local false discovery rate and agreement with covariation.** **a**, Right side of the scan distribution (black points), the mixture density fitted by Lindsey's method (blue) and the empirical null $\pi_0 f_0$ (dashed). **b**, Local false discovery rate lfdr $= \pi_0 f_0(z)/f(z)$, made monotonically non-increasing on the right: 0.93 at the top-1% boundary and 0.13 at the top 0.1%; lfdr ≤ 0.2 corresponds to z ≥ 2.72 (P ≤ 0.0033). The background fraction of the top 1% recomputed from the lfdr agrees with Efron's tail-area estimate (0.612 versus 0.613). **c**, Fraction of candidates with covariation in the structure-free alignment, by R-scape power class (expected number of covarying base pairs), for candidates with statistical support (lfdr ≤ 0.2 or a BH window; 4,372 candidates with an evaluable alignment) and without it (11,438), with Wilson 95% intervals (7.3% versus 3.6% overall; two-sided Fisher's exact P = 2 × $10^{-21}$). Although the lfdr is computed from the scan score alone, covariation is more frequent at candidates with low lfdr, and the difference persists in all four power classes (two-sided Fisher's exact P ≤ 0.03 in each, not adjusted for multiple comparisons).

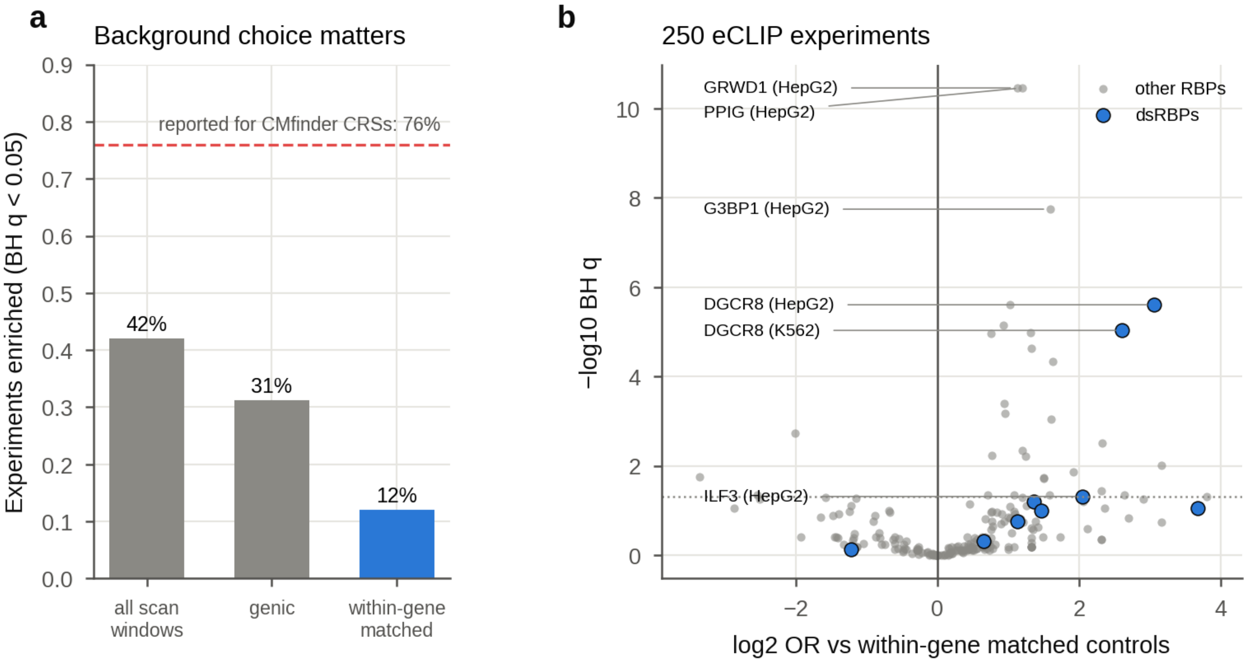


**Supplementary Fig. S9 | Design of the eCLIP analysis: choice of background and enrichment per experiment. a**, Fraction of the 250 eCLIP experiments enriched at candidates at BH $q < 0.05$: 42% against all scan windows, 31% against windows in transcribed regions and 12% against within-gene matched controls. Against scan windows, the tests are one-sided binomial z-tests within ten GC bins, as in Seemann 2017; against matched controls, two-sided Cochran–Mantel–Haenszel tests with each candidate and its controls as one stratum; BH adjustment is across the 250 experiments. The dashed red line marks the value reported by Seemann 2017 for the CMfinder CRSs (76%). **b**, Enrichment per experiment (Mantel–Haenszel log2 odds ratio against same-gene controls and BH q); blue, dsRBPs. The dsRBPs together (candidates bound by any of them; Mantel–Haenszel; 15,245 candidates and 73,137 controls): odds ratio 2.2 (95% interval 1.5–3.2), and 1.5 (two-sided Cochran–Mantel–Haenszel $P = 0.06$; 15,184 candidates) without candidates near known miRNA genes; DGCR8 alone, odds ratio 8.3 in HepG2 and 6.1 in K562.

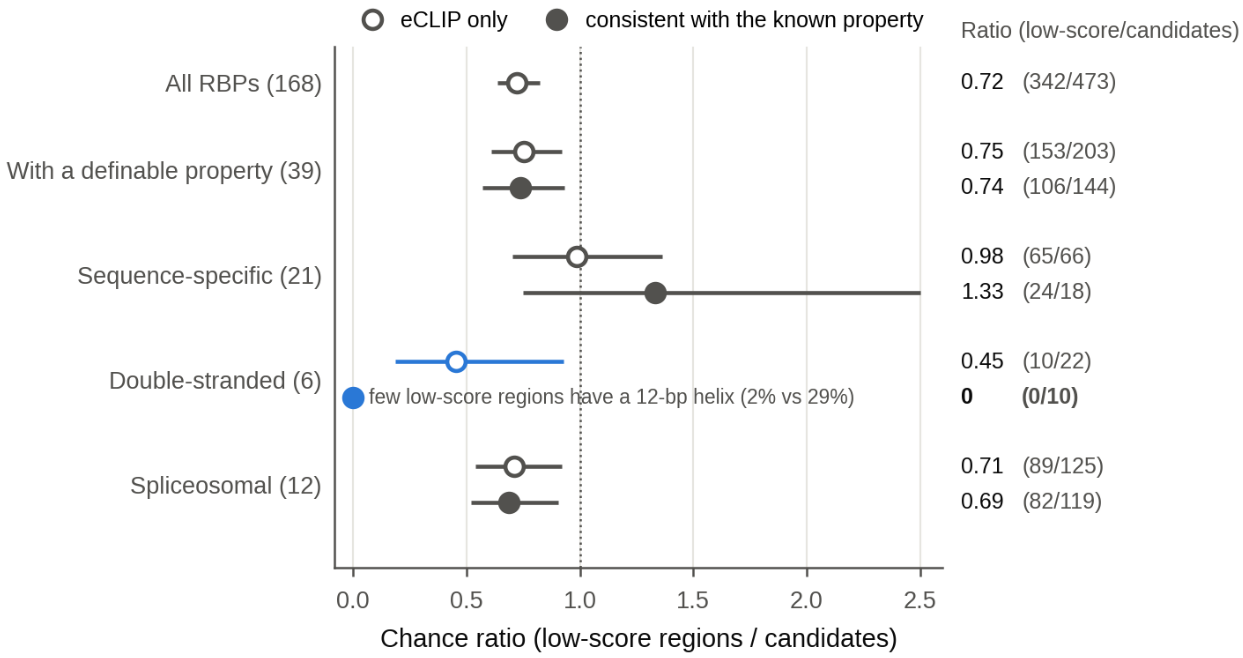


**Supplementary Fig. S10 | Chance ratio when binding calls are restricted to those consistent with the partner's known properties.** Ratio of the numbers of low-score regions and candidates with a candidate-specific RBP (chance ratio), with 95% intervals from 2,000 bootstrap resamples of the 6,088 pairs of a candidate and a low-score region (Fig. 6b). Open circles, eCLIP calls only; filled circles, only calls consistent with the partner's known property. The column on the right gives the ratio and the counts (low-score regions / candidates). With eCLIP calls only, 0.72 for all 168 RBPs (342 / 473). For the 39 RBPs with a definable property, 0.75 with eCLIP only (153 / 203) and 0.74 when restricted to calls consistent with the property (106 / 144). For the 21 sequence-specific RBPs (recognition sequence, the top 10 enriched 5-mers from ENCODE RNA Bind-n-Seq, Dominguez 2018; property, the recognition sequence lies in a single-stranded part), the ratio worsened from 0.98 to 1.33. For the dsRBPs (6, blue; helix of at least 12 bp in the MFE structure), it fell from 0.45 (95% interval 0.19–0.93) to 0 (10 candidates, 0 low-score regions); because such helices occur in about 29% of candidates but about 2% of low-score regions, this fall is expected even without binding specificity and is not evidence of it. The worse ratio for sequence-specific RBPs is the converse: their property requires unpaired bases, which favours the less structured low-score regions. For spliceosomal factors (12; within 100 nt of a splice site), it went from 0.71 to 0.69. The dotted line marks a ratio of 1.

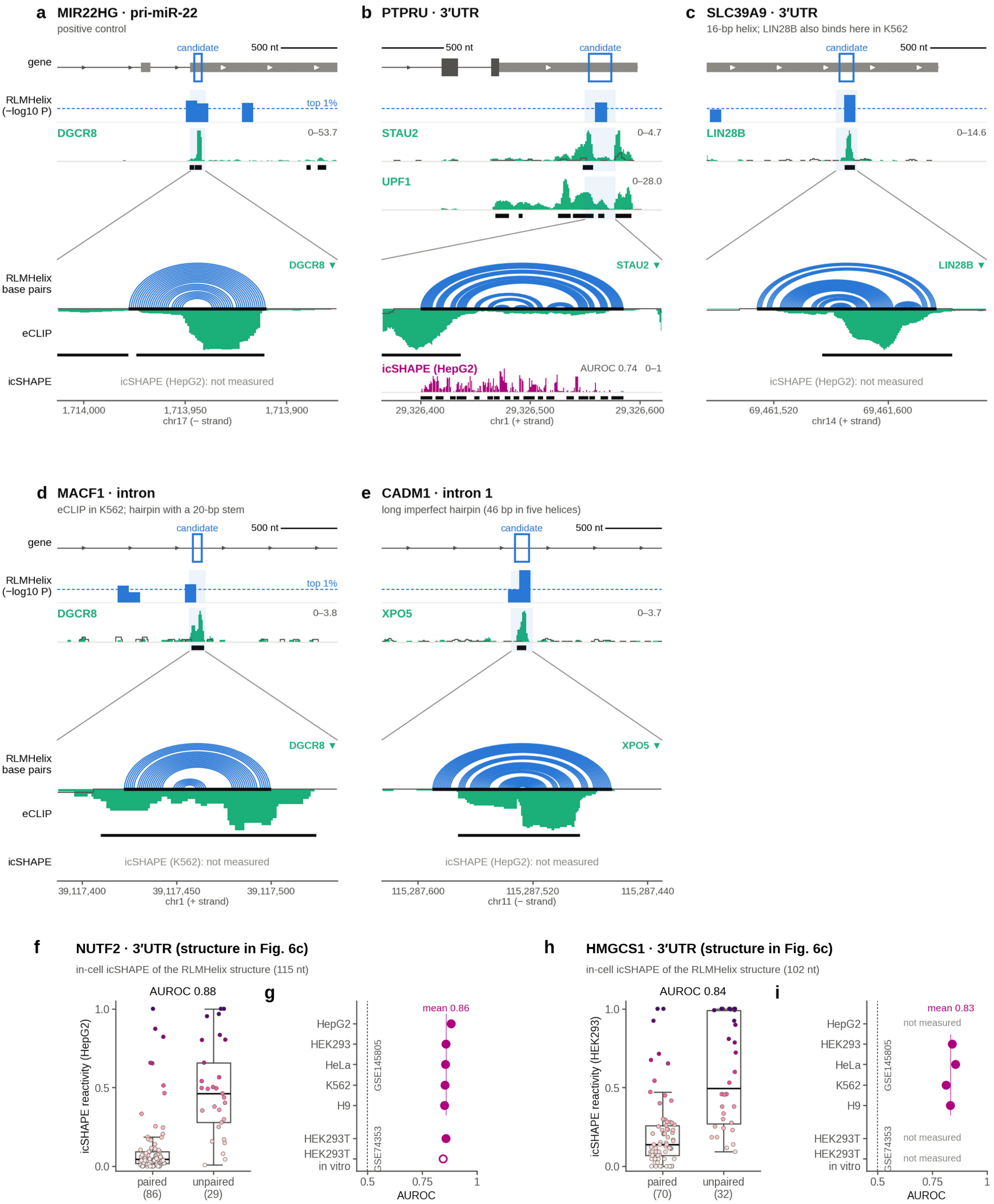

a MIR22HG · pri-miR-22
positive control
b PTPRU · 3′UTR
c SLC39A9 · 3′UTR
16-bp helix; LIN28B also binds here in K562
gene
RLMHelix (−log10 P)
candidate
500 nt
top 1%
DGCR8
STAU2
UPF1
LIN28B
RLMHelix base pairs
eCLIP
icSHAPE
icSHAPE (HepG2): not measured
icSHAPE (HepG2)
AUROC 0.74
chr17 (− strand)
chr1 (+ strand)
chr14 (+ strand)
d MACF1 · intron
eCLIP in K562; hairpin with a 20-bp stem
e CADM1 · intron 1
long imperfect hairpin (46 bp in five helices)
XPO5
icSHAPE (K562): not measured
chr11 (− strand)
f NUTF2 · 3′UTR (structure in Fig. 6c)
in-cell icSHAPE of the RLMHelix structure (115 nt)
AUROC 0.88
icSHAPE reactivity (HepG2)
paired (86)
unpaired (29)
g
mean 0.86
HepG2
HEK293
HeLa
K562
H9
HEK293T
HEK293T in vitro
GSE145805
GSE74353
AUROC
h HMGCS1 · 3′UTR (structure in Fig. 6c)
in-cell icSHAPE of the RLMHelix structure (102 nt)
AUROC 0.84
icSHAPE reactivity (HEK293)
paired (70)
unpaired (32)
i
mean 0.83
not measured

**Supplementary Fig. S11 | eCLIP and icSHAPE at five more candidates, and in-cell icSHAPE on the *NUTF2* and *HMGCS1* 3′UTR structures.** **a**–**e**, Five candidates drawn as in Fig. 6c (ENCODE eCLIP in HepG2; in K562 for **d**), with in-cell (in vivo) icSHAPE reactivity added under each zoom (GSE145805, in the cell line of the eCLIP: HepG2, or K562 for **d**; the source study's processed icSHAPE-pipe scores, 0–1, used as provided); black bars under the icSHAPE track, nucleotides paired in the RLMHelix structure. **a**, The pri-miR-22 positive control (RLMFam03356; DGCR8): a sharp DGCR8 signal of the Microprocessor lies on the predicted 15-base-pair helix. **b**, The *PTPRU* 3′UTR (RLMFam03987; STAU2 and UPF1): the STAU2 signal, absent from the input, rises just outside the structure on both sides, with its peak on the 5′ arm of an 11-base-pair helix, and UPF1 binds the whole 3′UTR broadly. icSHAPE reactivity is lower over the predicted helices than in the loops (AUROC 0.74 in HepG2; icSHAPE agreement, the mean of four cell lines, 0.76). **c**–**e**, Candidates in which a structure-recognizing RBP binds the predicted helix: the *SLC39A9* 3′UTR (RLMFam02146; LIN28B, which also binds here in K562), an intron of *MACF1* (RLMFam04531; DGCR8 in K562, on a hairpin with a 20-base-pair stem; no annotated miRNA nearby) and the first intron of the main *CADM1* transcript (5′UTR of a minor transcript; RLMFam00511; exportin-5, XPO5, on a long imperfect hairpin of 46 base pairs in five helices). None of these three cores, nor the pri-miR-22 core, has a measured nucleotide in the icSHAPE data of the cell line drawn, so their icSHAPE rows read "not measured". **f**–**i**, The two 3′UTR candidates of Fig. 6c (their structures are drawn in Fig. 6c). **f**, **g**, *NUTF2* (RLMFam00092). **f**, Reactivity of the 86 paired and 29 unpaired nucleotides in HepG2 (boxes, median and quartiles; whiskers, the most extreme values within 1.5 × the interquartile range; points, nucleotides); AUROC of reactivity for unpaired versus paired nucleotides, 0.88. **g**, AUROC in each of the five cell lines of GSE145805 (0.85–0.88; their mean, 0.86, is the icSHAPE agreement given in the text) and in HEK293T of GSE74353 (in cell, 0.86; open circle, in vitro, 0.84). **h**, **i**, *HMGCS1* (RLMFam11457). **h**, Reactivity of the 70 paired and 32 unpaired nucleotides in HEK293 (AUROC 0.84). **i**, AUROC in the four GSE145805 cell lines that measured the core (0.81–0.85; mean 0.83, the icSHAPE agreement in the text); in HepG2 and in HEK293T of GSE74353, less than half of the core was measured (not measured). Dashed lines, 0.5. For both candidates and in every cell line, the AUROC was higher than with any circular shift of the paired/unpaired labels (114 shifts for *NUTF2*, 101 for *HMGCS1*; one-sided empirical P = 0.009 and 0.010, computed as (1 + the number of shifts with an AUROC at least as high)/(1 + the number of shifts)).

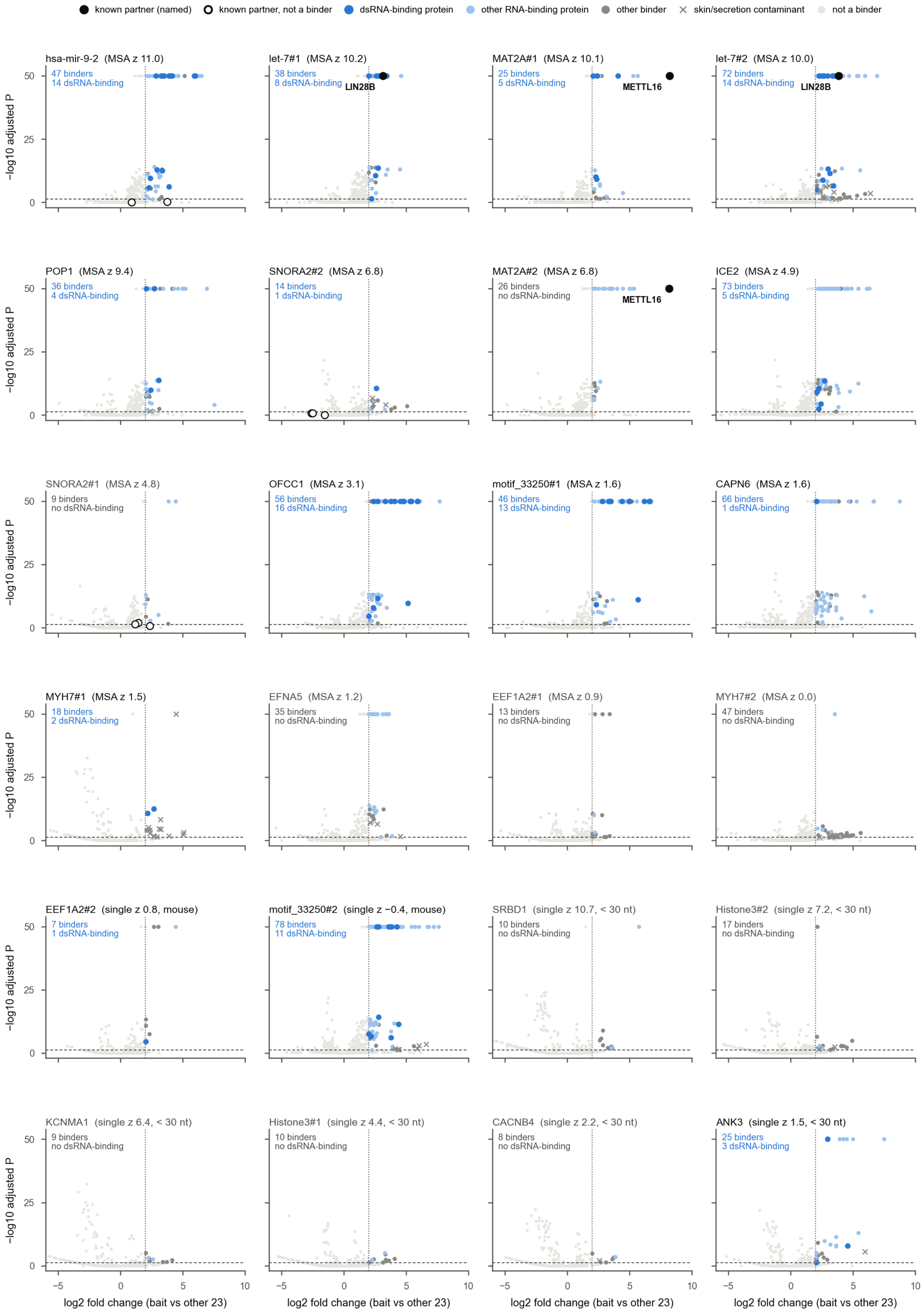

known partner (named)
known partner, not a binder
dsRNA-binding protein
other RNA-binding protein
other binder
skin/secretion contaminant
not a binder
hsa-mir-9-2 (MSA z 11.0)
47 binders
14 dsRNA-binding
let-7#1 (MSA z 10.2)
38 binders
8 dsRNA-binding
LIN28B
MAT2A#1 (MSA z 10.1)
25 binders
5 dsRNA-binding
METTL16
let-7#2 (MSA z 10.0)
72 binders
14 dsRNA-binding
LIN28B
POP1 (MSA z 9.4)
36 binders
4 dsRNA-binding
SNORA2#2 (MSA z 6.8)
14 binders
1 dsRNA-binding
MAT2A#2 (MSA z 6.8)
26 binders
no dsRNA-binding
METTL16
ICE2 (MSA z 4.9)
73 binders
5 dsRNA-binding
SNORA2#1 (MSA z 4.8)
9 binders
no dsRNA-binding
OFCC1 (MSA z 3.1)
56 binders
16 dsRNA-binding
motif_33250#1 (MSA z 1.6)
46 binders
13 dsRNA-binding
CAPN6 (MSA z 1.6)
66 binders
1 dsRNA-binding
MYH7#1 (MSA z 1.5)
18 binders
2 dsRNA-binding
EFNA5 (MSA z 1.2)
35 binders
no dsRNA-binding
EEF1A2#1 (MSA z 0.9)
13 binders
no dsRNA-binding
MYH7#2 (MSA z 0.0)
47 binders
no dsRNA-binding
EEF1A2#2 (single z 0.8, mouse)
7 binders
1 dsRNA-binding
motif_33250#2 (single z −0.4, mouse)
78 binders
11 dsRNA-binding
SRBD1 (single z 10.7, < 30 nt)
10 binders
no dsRNA-binding
Histone3#2 (single z 7.2, < 30 nt)
17 binders
no dsRNA-binding
KCNMA1 (single z 6.4, < 30 nt)
9 binders
no dsRNA-binding
Histone3#1 (single z 4.4, < 30 nt)
10 binders
no dsRNA-binding
CACNB4 (single z 2.2, < 30 nt)
8 binders
no dsRNA-binding
ANK3 (single z 1.5, < 30 nt)
25 binders
3 dsRNA-binding
−log10 adjusted P
0
25
50
−5
0
5
10
log2 fold change (bait vs other 23)

**Supplementary Fig. S12 | Binders recovered by the pull-down mass spectrometry.** Volcano plots of all 24 RNA baits. Each bait was compared with the mean of the other 23 by a two-sided Wald test in proDA (log2-transformed, median-normalized intensities; four runs per bait, two pull-downs each measured twice), and P values were adjusted by Benjamini–Hochberg across proteins for each bait. x, log2 fold change; y, adjusted P (−log10, capped at 50). Proteins with adjusted $P < 0.05$ (dashed line) and log2 fold change ≥ 2 (dotted line) are binders. Dark blue, double-stranded RNA-binding proteins (fixed list and Pfam double-stranded RNA-binding domains); light blue, other RNA-binding proteins; grey, other binders. Black, known partners called as binders (named: LIN28B for both let-7 baits, METTL16 for both *MAT2A* hairpins); open circles, known partners not called. Histone3#2 follows the strand of the EvoFold prediction, the reverse complement of the histone stem-loop bound by SLBP, so SLBP binding is not expected for this bait. Crosses, typical skin and secretion contaminants (61 proteins such as keratins, S100A7/8/9, SERPINB3/4, DCD and LCN1, which were kept in the data, the binder counts and the labels and only drawn with a separate symbol). Top left, number of binders (counted per protein group) and of double-stranded RNA-binding proteins among them. Order: the 16 baits analysed in Fig. 6f (by RLMHelix-MSA z), the 2 mouse baits and the 6 baits shorter than 30 nt (by single-sequence z). Baits with grey titles have no structure-recognizing binder.

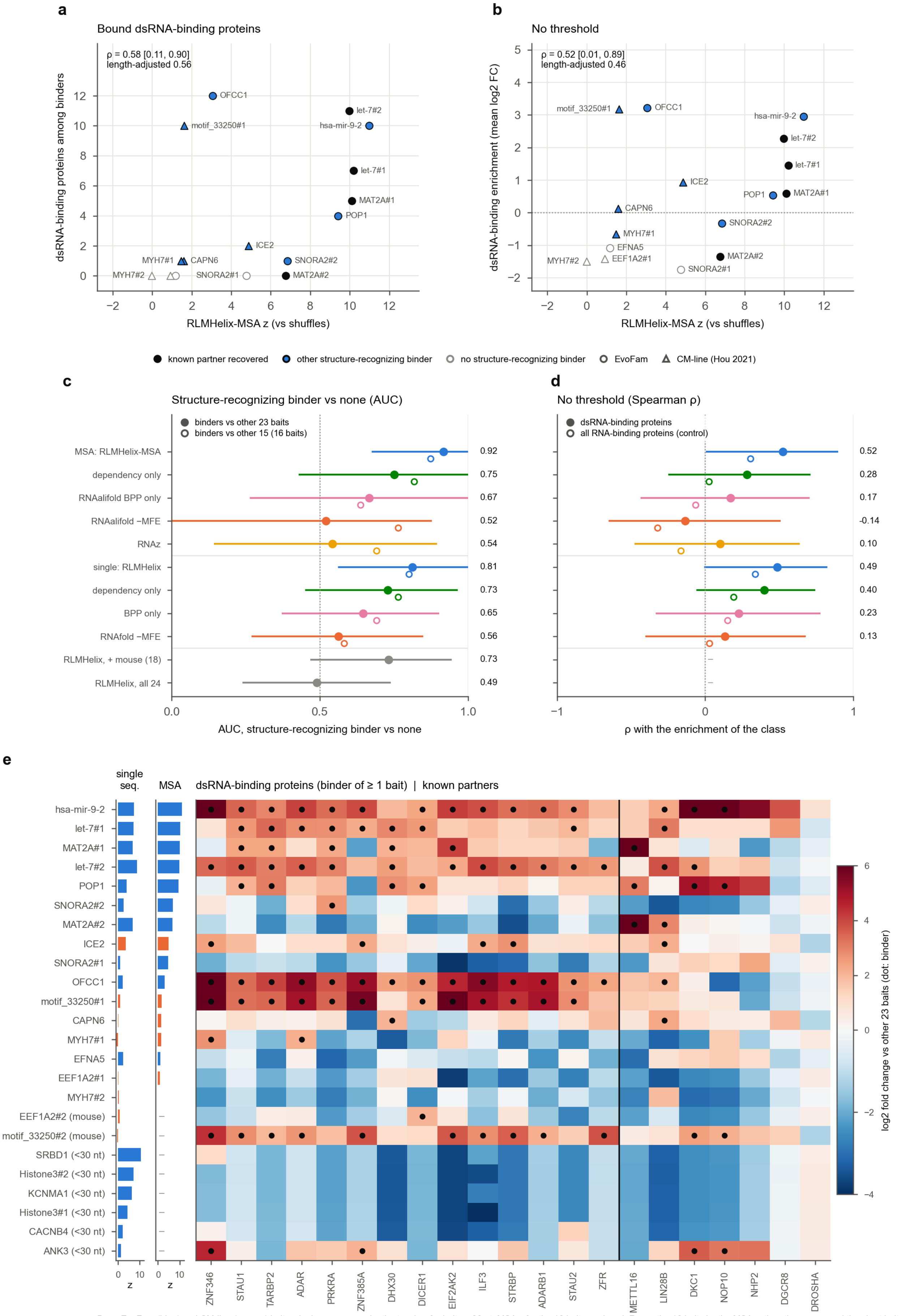


Bars: EvoFam (blue) and CM-line (orange) baits; single-sequence z by the teacher for baits < 30 nt; MSA z for the 16 baits analysed. Rows: the 16 baits by the MSA z, then the mouse and the short baits.

**Supplementary Fig. S13 | In the pull-downs, recruitment of double-stranded RNA-binding proteins tracks the RLMHelix-MSA score (exploratory).** Binders were called with all 24 baits, each against the other 23 (Supplementary Fig. S12, Supplementary Table S7). **a**, Shuffle z of RLMHelix-MSA against the number of bound double-stranded RNA-binding proteins of the fixed list of 17 (16 human baits of ≥30 nt; Supplementary Methods; the counts in Supplementary Fig. S12 also include proteins with a Pfam double-stranded RNA-binding domain); Spearman $\varrho$ = 0.58 (bootstrap 95% interval 0.11–0.90; two-sided P = 0.017; length-adjusted 0.56). **b**, z against the mean log2 fold change of all quantified double-stranded RNA-binding proteins (fixed list and Pfam double-stranded RNA-binding domains, as in Supplementary Fig. S12): $\varrho$ = 0.52 (0.01–0.89; P = 0.037; length-adjusted 0.46). In **a** and **b**: black, known partner recovered; blue, another structure-recognizing binder; white, none; circles, EvoFam; triangles, CM-line. **c**, AUC (95% interval) of each score's z for baits with versus without a structure-recognizing binder: five alignment and four single-sequence scores (16 baits), single-sequence RLMHelix with the two mouse baits (18 baits; 0.73) and with the six baits shorter than 30 nt (24 baits, teacher; 0.49). Open symbols, binders called among the 16 baits only (RLMHelix-MSA 0.87, single sequence 0.80). For single-sequence RLMHelix on the 16 baits, AUC 0.81 (0.56–1.00), two-sided Mann–Whitney P = 0.078 and two-sided paired DeLong P = 0.020 against RNAfold −MFE. **d**, Spearman $\varrho$ of the comparison in **b** for each score (RLMHelix-MSA 0.52, dependency only 0.28, RNAalifold −MFE −0.14, RNAz 0.10; RLMHelix 0.49, RNAfold −MFE 0.13); open symbols, the same over all RNA-binding proteins (RLMHelix-MSA 0.31, RLMHelix 0.34, RNAalifold −MFE −0.32, RNAz −0.16). **e**, Log2 fold change of double-stranded RNA-binding proteins (fixed list and Pfam domains) and known partners (columns; dots, binders) for the 24 baits (rows); bars, z of single-sequence RLMHelix and RLMHelix-MSA (blue, EvoFam; orange, CM-line). Short baits bind few double-stranded RNA-binding proteins even when their single-sequence scores are high. Intervals are 95% percentile intervals from 10,000 bootstrap resamples of baits; P values are not adjusted for multiple comparisons.

### Supplementary tables

**Supplementary Table S1.** Evaluation sets and denominators (Rfam 328/322, EvoFam, RNAz 2005, scan windows)

**Supplementary Table S2.** Comparison with phylogenetic models (322 families)

**Supplementary Table S3.** Known functional structures (82 elements, PMIDs, how coordinates were determined)

**Supplementary Table S4.** RLM-Fam catalogue: one row per candidate (with its RLMFam ID; lfdr, evidence tier, selection route, annotation) and one row per family (RLMFam ID, as used in the text)

**Supplementary Table S5.** eCLIP results per experiment

**Supplementary Table S6.** Structure–partner pairings suggested by the analyses (21 entries, literature support, PMIDs)

**Supplementary Table S7.** Baits of the pull-down mass spectrometry (sequence, source, length, number of binders, known partners, empirical P values and z of each score; calls with all 24 baits and within the 16 baits)

**Supplementary Table S8.** Accessions of the public data used (eCLIP, control RNA-seq for the expression filter, RBNS, icSHAPE, bigWig tracks of the genome-browser views, PolyASite 2.0) and the 55 vertebrate genomes of the homology search (Ensembl release 116)
